# Declining ribosome function limits oocyte longevity

**DOI:** 10.64898/2026.09.29.755503

**Authors:** Kayla Judson, Katie J. Danielson, Shuang Tang, Sahil Chandhok, Elizabeth Hui, Fiona Wong, Thibault Mayor, Karen Schindler, Ethan J. Greenblatt

## Abstract

Protein synthesis declines with age across many tissues, but whether this decline actively contributes to physiological aging remains unclear. Oocytes may be particularly vulnerable because they rely on ribosomes synthesized during growth and maintained for prolonged periods without continual replacement. Here we show that declining translational capacity is a causal component of oocyte aging. An unbiased genetic screen identified ribosome dosage as a major determinant of oocyte longevity, and experimentally reducing ribosome abundance was sufficient to accelerate the loss of developmental competence and prematurely recapitulate meiotic defects that arise during natural aging, while also producing defects in early embryonic development. Ribosomes isolated from aged *Drosophila* oocytes exhibit ∼50% lower translational activity despite only a modest reduction in ribosome abundance and preserved gross structure and composition, with both ribosomal subunits contributing to this functional decline. Protein synthesis also declines markedly in aged metaphase-I mouse oocytes. Together, these findings identify long-lived translational machinery as an intrinsic vulnerability of the oocyte and establish declining ribosome function as a mechanistic contributor to reproductive aging.

## INTRODUCTION

Protein synthesis declines with age across diverse tissues and organisms (Danielson et al. 2024; Kim and Pickering 2023; Stein et al. 2022), but whether this decline actively drives functional aging or instead reflects deterioration elsewhere in the cell remains unresolved. Because translation is required to maintain the proteome, declining translational capacity could itself contribute to the progressive loss of cellular function with age. Oocytes provide an especially powerful system in which to test this possibility. With increasing maternal age, oocytes become more prone to meiotic chromosome segregation errors and lose developmental competence, limiting human reproductive lifespan in the years leading to menopause (Lister et al. 2010; Gruhn et al. 2019; Battaglia et al. 1996). Oocytes may be particularly vulnerable to declining translation because transcription largely ceases during late development, leaving meiotic progression and early embryonic development dependent on translation of stored maternal mRNAs (Susor et al. 2015; Sousa Martins et al. 2016). Moreover, ribosomes synthesized during oocyte growth are stockpiled for prolonged periods rather than continually replenished (De Leon et al. 1983), creating an unusual dependence on the long-term integrity of pre-existing translational machinery. Consistent with a link between ribosome insufficiency and reproductive longevity in humans, premature ovarian failure and increased miscarriage risk have been reported in Diamond–Blackfan anemia, a ribosomopathy caused predominantly by haploinsufficiency of ribosomal protein genes (Tufano et al. 2014; Leblanc et al. 2005). Together, these observations raise the possibility that deterioration of the translational machinery contributes directly to the age-dependent loss of oocyte function.

Several molecular changes have been implicated in the age-dependent loss of oocyte quality, although their causal contributions remain incompletely defined. Among the best characterized is the progressive loss of cohesin complexes that maintain sister chromatid cohesion. Because meiotic cohesin is established early in oocyte development, which occurs during fetal life, and must persist throughout reproductive life, its deterioration is thought to contribute substantially to age-related chromosome segregation errors (Tsutsumi et al. 2014; Chiang et al. 2010). However, cohesin loss can precede the pronounced increase in chromosome segregation defects (Chiang et al. 2010), indicating that additional age-dependent processes likely contribute to declining meiotic competence.

Determining which age-associated changes causally limit oocyte function is challenging in mammals because reproductive aging occurs over long timescales (i.e. decades), oocytes are available in limited quantities, and genetic manipulation is comparatively difficult. We therefore used *Drosophila melanogaster*, in which key features of oocyte development and meiotic regulation are conserved. Like mammalian oocytes, mature *Drosophila* oocytes undergo a prolonged meiotic arrest, although at metaphase I rather than metaphase II. *Drosophila* females produce approximately 100 mature oocytes per day, providing sufficient material for biochemical analysis while allowing precise control of oocyte age and extensive genetic manipulation. Oocyte viability can also be followed quantitatively using hatch-rate curves, which decline reproducibly with increasing storage time (Greenblatt et al. 2019). Importantly, this decline is largely oocyte autonomous, allowing age-dependent changes intrinsic to the oocyte to be distinguished from systemic aging of the animal.

*Drosophila* oocytes develop aging-associated metaphase I spindle defects resembling those observed in aged mammalian oocytes, including loss of spindle bipolarity (Holubcova et al. 2015; Nakagawa and FitzHarris 2017; Greenblatt et al. 2019). Notably, these defects arise over the same period in which protein synthesis declines by approximately 50% in aging *Drosophila* oocytes (Greenblatt et al. 2019). In mammalian oocytes, age-dependent translational changes affect mRNAs associated with meiotic and spindle function (Del Llano et al. 2020), and the abundance or localization of several proteins required for meiotic spindle function declines with maternal age (Yun et al. 2014; Lagirand-Cantaloube et al. 2017; Zielinska et al. 2019; Yin et al. 2023). Together, these observations raise the possibility that declining translational capacity compromises the production of proteins required for meiotic spindle function during oocyte aging.

Here, we combine genetic and biochemical approaches to investigate the basis and functional consequences of declining translation during oocyte aging. We show that ribosomes isolated from aged *Drosophila* oocytes have substantially reduced translational activity despite no detectable large-scale changes in ribosomal protein composition or rRNA structure, with both the small and large ribosomal subunits contributing to this loss of activity. We further show that oocytes are highly sensitive to reductions in ribosome abundance: genetically reducing ribosome content accelerates the loss of developmental competence and produces premature meiotic spindle defects resembling those that arise during natural oocyte aging. Finally, we find that protein synthesis also declines markedly with age in metaphase I mouse oocytes, indicating that age-associated translational decline extends across species. Together, our findings identify declining ribosome function as a causal contributor to the age-dependent loss of oocyte quality and provide a mechanistic link between translational decline and reproductive aging.

## RESULTS

### Oocyte longevity is highly sensitive to ribosome content

We sought to determine whether an unbiased genetic approach would identify cellular pathways that become functionally limiting during oocyte aging. We reasoned that reducing gene dosage by half could expose processes that are well tolerated in young oocytes but become limiting as oocytes age and cellular function declines. Such genes would therefore be expected to have little or no effect on young oocytes, yet cause a premature loss of viability following prolonged storage. This sensitized strategy also allowed us to interrogate essential genes that could not readily be assessed using homozygous loss-of-function mutations. We therefore performed an unbiased genetic screen using *Drosophila* deficiency lines heterozygous for deletions spanning multiple genes, covering the majority of chromosome 2, with hatch rate as a quantitative measure of oocyte viability. Deficiency lines were outcrossed with wild-type strains to minimize effects of background mutations, and hatch rates were compared between young, unstored oocytes and oocytes aged for 10 days, a timepoint at which control oocytes show little decline in viability (Fig. 1A). We focused on deficiency lines that retained high viability when young but exhibited a pronounced reduction in hatch rate following storage, thereby identifying genomic regions whose reduced dosage selectively impaired the maintenance of oocyte function over time rather than causing a constitutive developmental defect. Overlapping deficiencies were then used to narrow candidate regions, with genes not detectably expressed or translated in mature oocytes excluded using existing RNA-seq and ribosome-profiling datasets (Brown et al. 2014; Graveley et al. 2011; Greenblatt et al. 2019) (Fig. 1B).

**Figure 1.**
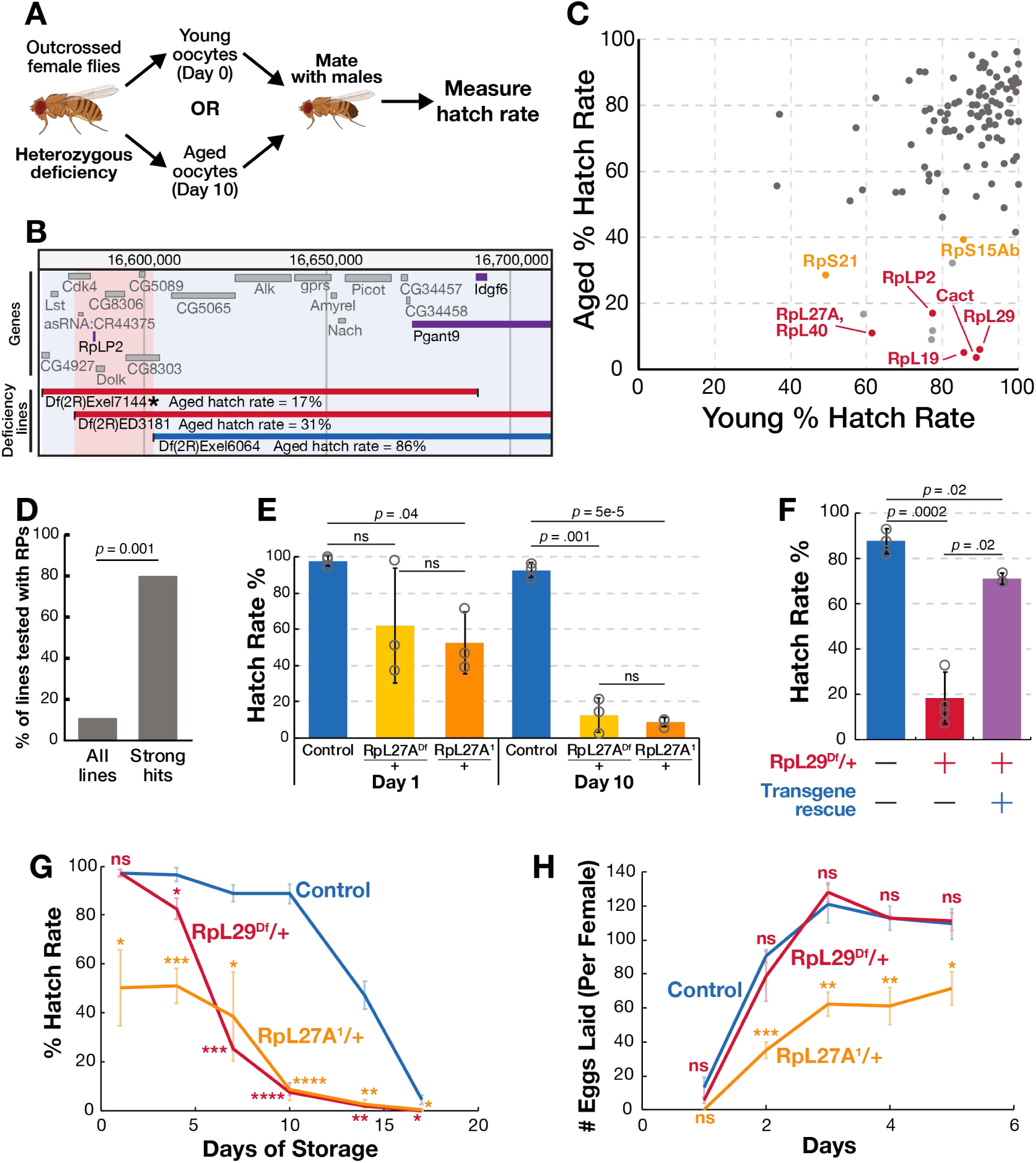
An unbiased genetic screen identifies ribosomal protein genes as prominent dosage-sensitive determinants of oocyte longevity. **(A)** Schematic of the genetic screen design used to identify dosage-sensitive pathways in aged but not young oocytes. **(B)** Schematic showing the use of overlapping deficiency lines to narrow down candidate genes from the genetic screen. Genes labeled in purple and grey are considered expressed or not expressed respectively. (C) Plot summarizing the screen results. Each point represents results obtained from a single deficiency line: red, strong hits; yellow, moderate hits. Deficiency lines where a single candidate gene could not be resolved through mapping experiments are shown in light grey. **(D)** Ribosomal protein (RP) genes are strongly enriched among strong hits relative to their frequency among all deficiency lines screened, Fisher’s exact test is reported. **(E)** Oocytes heterozygous for the *RpL27A^1^*mutant allele show similar hatch rates after 1 and 10 days as compared to the *RpL27A^Df^* deficiency line. **(F)** Hatch rate of heterozygous deficiency line, *RpL29^Df^*/+ oocytes after 10 days of oocyte storage is strongly rescued by germline-specific expression of *RpL29*. **(G)** Hatch rate curves over storage time show accelerated decline in oocyte viability for both *RpL29^Df^*/+ and *RpL27A^1^*/+ flies. **(H)** Egg-laying rate per female per day shows no significant change in *RpL29^Df^*/+ animals as compared to wild type animals, whereas heterozygous *RpL27A^1^* flies lay approximately half as many eggs. Welch’s unpaired *t*-test, \**P* < 0.05, \*\**P* < 0.005, \*\*\**P* < 0.0005, \*\*\*\**P* < 0.00005.

The screen encompassed more than 110 deficiency lines covering over 4,000 genes, yet relatively few regions produced a strong age-dependent loss of viability (Fig. 1C, Supplementary Table S1). Three hits could not be resolved to a single gene by overlapping deficiency mapping and were not pursued further (Fig. S1). Strikingly, four of the five strongest hits contained ribosomal protein (RP) genes, whereas the remaining hit contained the developmental patterning gene *cactus*. RP-containing deficiencies were strongly enriched among the strongest hits: only 11% of all lines screened contained an RP gene, compared with 80% of the strongest hits (Fig. 1D, *p* = 0.001). We therefore examined additional deficiencies spanning RP genes and found that most reduced hatch rates from aged oocytes, whereas deficiencies spanning key meiotic genes had only modest effects (Table 1). Thus, the unbiased genetic screen independently identified ribosome dosage as a major determinant of oocyte longevity.

**Table 1.** Ribosomal protein and spindle-associated genes covered in the genetic screen, with corresponding hatch rate and ribosome profiling TPM.

|  |  | Genes | Stg 14 Expression (TPM) | D0 Hatch Rate % | D10 Hatch Rate % |
| --- | --- | --- | --- | --- | --- |
| Ribosomal Genes | Ribosomal proteins | RpL41 | 11747 | 95 | 87 |
|  |  | RpS18 | 2906 | 81 | 76 |
|  |  | RpL27A* | 2286 | 62 | 13 |
|  |  | RpL38 | 2156 | 80 | 46 |
|  |  | RpS24** | 1987 | 76 | 59 |
|  |  | RpLP2 | 1918 | 73 | 17 |
|  |  | RpL40* | 1432 | 62 | 13 |
|  |  | RpL29 | 1174 | 90 | 6 |
|  |  | RpS16** | 954 | 76 | 59 |
|  |  | RpL19 | 533 | 86 | 6 |
|  |  | RpS15Ab | 527 | 86 | 40 |
|  |  | RpS21 | 511 | 49 | 30 |
|  |  | RpS2 | 498 | 94 | 77 |
| Meiotic Genes | Spindle-associated genes | Eb1 | 185 | 97 | 78 |
|  |  | sub | 126 | 94 | 87 |
|  |  | BubR1 | 115 | 91 | 92 |
|  |  | cid | 91 | 90 | 93 |
|  |  | Mad1 | 69 | 88 | 88 |
|  |  | SMC1 | 58 | 71 | 70 |
|  |  | mars | 57 | 90 | 93 |
|  |  | Rcc1 | 29 | 33 | 77 |
|  |  | mei-S332 | 15 | 62 | 82 |
|  |  | Patronin | 4 | 90 | 83 |

\*, \*\* mark genes covered in the same deficiency line

To identify the individual genes responsible for the age-dependent phenotypes observed in the deficiency screen, we tested partially overlapping deficiencies, analyzed available mutant alleles, and performed genetic rescue experiments. For *RpL27A*, an independent *RpL27A¹/+* mutant exhibited an age-dependent reduction in hatch rate similar to that of a deficiency spanning *RpL27A*, identifying reduced *RpL27A* dosage as responsible for the phenotype (Fig. 1E). Because no independent alleles were available for another strong hit containing *RpL29*, we performed cell type-specific rescue experiments. Using the germline-specific nanos-Gal4 driver, we restored *RpL29* expression in the germline of flies heterozygous for the *RpL29*-containing deficiency. Germline-specific expression of *RpL29* largely restored the hatch rate of aged oocytes from ∼19% to ∼71% (Fig. 1F). Together, these experiments identify individual RP genes underlying the deficiency phenotypes and demonstrate that reduced *RpL29* dosage in the germline is responsible for much of the premature decline in oocyte viability.

We next measured hatch rate over time to characterize oocyte longevity. Hatch rates fell to 50% after 5 days of storage in *RpL27A¹/+* animals and after 6 days in *RpL29^Df^*/+, roughly 8–9 days earlier than in controls (Fig. 1G). We prioritized *RpL29^Df^*/+ for further study because, unlike *RpL27A¹/+*, it showed no defects in young animals: hatch rate, egg production, and developmental timing were all comparable to controls (Fig. 1G,H). In contrast, *RpL27A¹/+* showed reduced hatch rate (51% vs. 97% in controls) and egg laying (66 vs. >100 eggs per fly per day). Consistent with this lack of baseline defects, *RpL29* is not classified as a Minute gene, a class of ribosomal protein genes whose reduced dosage produces dominant developmental phenotypes (Marygold et al. 2007). Finally, the *RpL29* deficiency is relatively small, covering 14 genes with only four oocyte-expressed genes compared with an average of 58 genes removed across the hit deficiency lines, making it well suited for studying the effects of reduced ribosome abundance on oocyte aging.

We next asked how loss of one copy of an RP affects oocyte ribosome abundance and composition. Using total 18S and 28S rRNA as a proxy for overall ribosome abundance, we observed reductions of 17% and 35% in *RpS21^Df^*/+ and *RpL29^Df^*/+ oocytes, respectively (Fig. 2A). Notably, *RpL29* loss reduced both 18S and 28S rRNA, whereas *RpS21* loss only reduced 18S rRNA (Fig. S2). This asymmetry may reflect the requirement for mature 60S subunits during late 40S maturation (Robledo et al. 2008; Lebaron et al. 2012). Proteomic analysis revealed no detectable changes in the composition of either ribosomal subunit (Fig. 2B), consistent with previous observations that RP loss primarily reduces ribosome biogenesis rather than altering the composition of mature ribosomes (Khajuria et al. 2018). The broader effect of *RpL29* hemizygosity on both ribosomal subunits could also contribute to the stronger effects of large-subunit RP deficiencies on aged hatch rate observed in our genetic screen (Table 1). Together, these results indicate that relatively modest reductions in overall ribosome abundance can substantially impair oocyte longevity.

**Figure 2.**
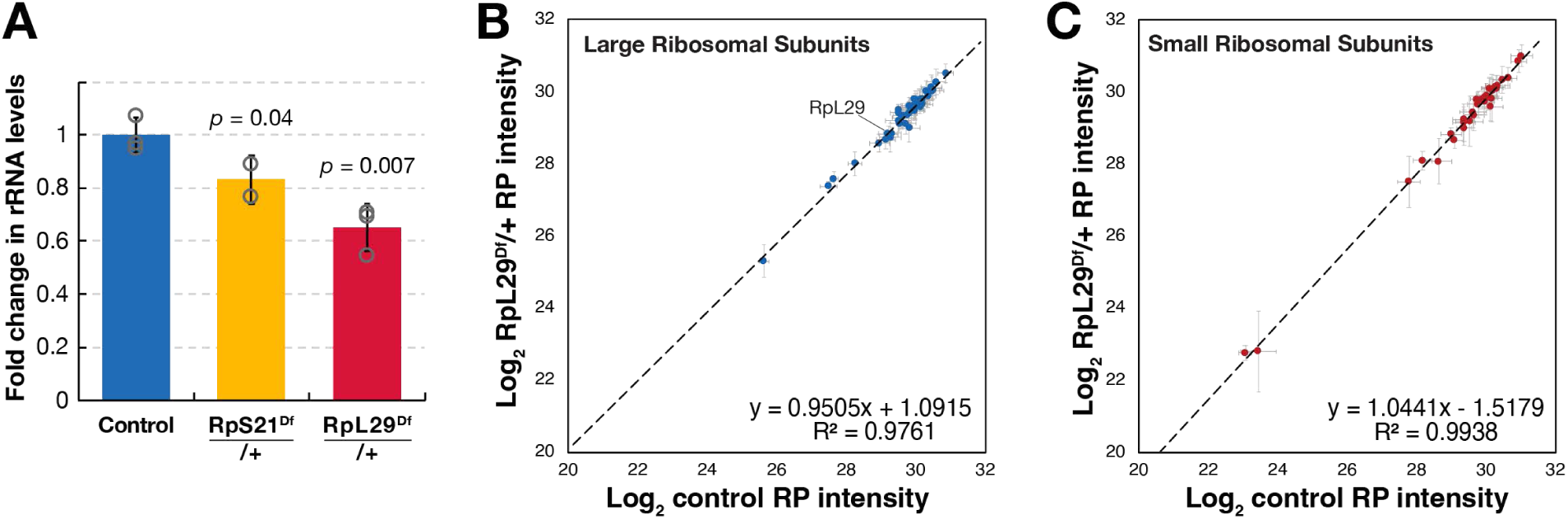
RP mutants reduce oocyte ribosome content but not ribosome composition. **(A)** Spike-in normalized total rRNA levels (18S + 28S) decrease by 17% and 35% in *RpS21^Df^*/+ and *RpL29^Df^*/+ flies respectively relative to wild type controls. Welch’s unpaired *t*-tests are reported. **(B-C)** Ribosomal protein composition is unchanged in *RpL29^Df^*/+ flies. Log_2_ intensity of ribosomal proteins in day 2 control (OrR) versus day 2 *RpL29^Df^*/+ by mass spectrometry. Pearson correlation coefficients are reported. N = 4 biological replicates per condition. **(B)** Large ribosomal subunits are shown in blue. RpL29 is highlighted to show no observable change in its stoichiometry. **(C)** Small ribosomal subunits are shown in red.

### Intrinsic ribosome activity decreases in aged oocytes

We next sought to test whether the sensitivity of aged oocytes to reduced ribosome dosage was reflective of an overall decline in their intrinsic ribosome activity. Using an established protocol for collecting *Drosophila* oocytes of defined ages (Greenblatt et al. 2019), we isolated young (2-day-old) and aged (14-day-old) oocytes, whose ability to support embryonic development to hatching declines from ∼98% to ∼50%, respectively. To directly measure ribosome activity, we developed an *in vitro* translation assay in which ribosomes isolated from young or aged oocytes were added to identical ribosome-depleted cytosolic extracts prepared from young oocytes (Fig. 3A). This design attributed differences in translation to the ribosomes themselves, rather than to soluble translation factors, tRNAs, or metabolites. To further minimize contributions from ribosome-associated initiation factors, we used a luciferase reporter driven by the cricket paralysis virus (CrPV) intergenic region internal ribosome entry site (IGR IRES), which initiates translation independently of canonical initiation factors (Wilson et al. 2000). Under these conditions, ribosomes isolated from aged oocytes exhibited approximately 50% lower translational activity than ribosomes isolated from young oocytes (Fig. 3B,C). This reduction was reproducible across three independent biological replicates and was observed across reporter mRNA concentrations.

**Figure 3.**
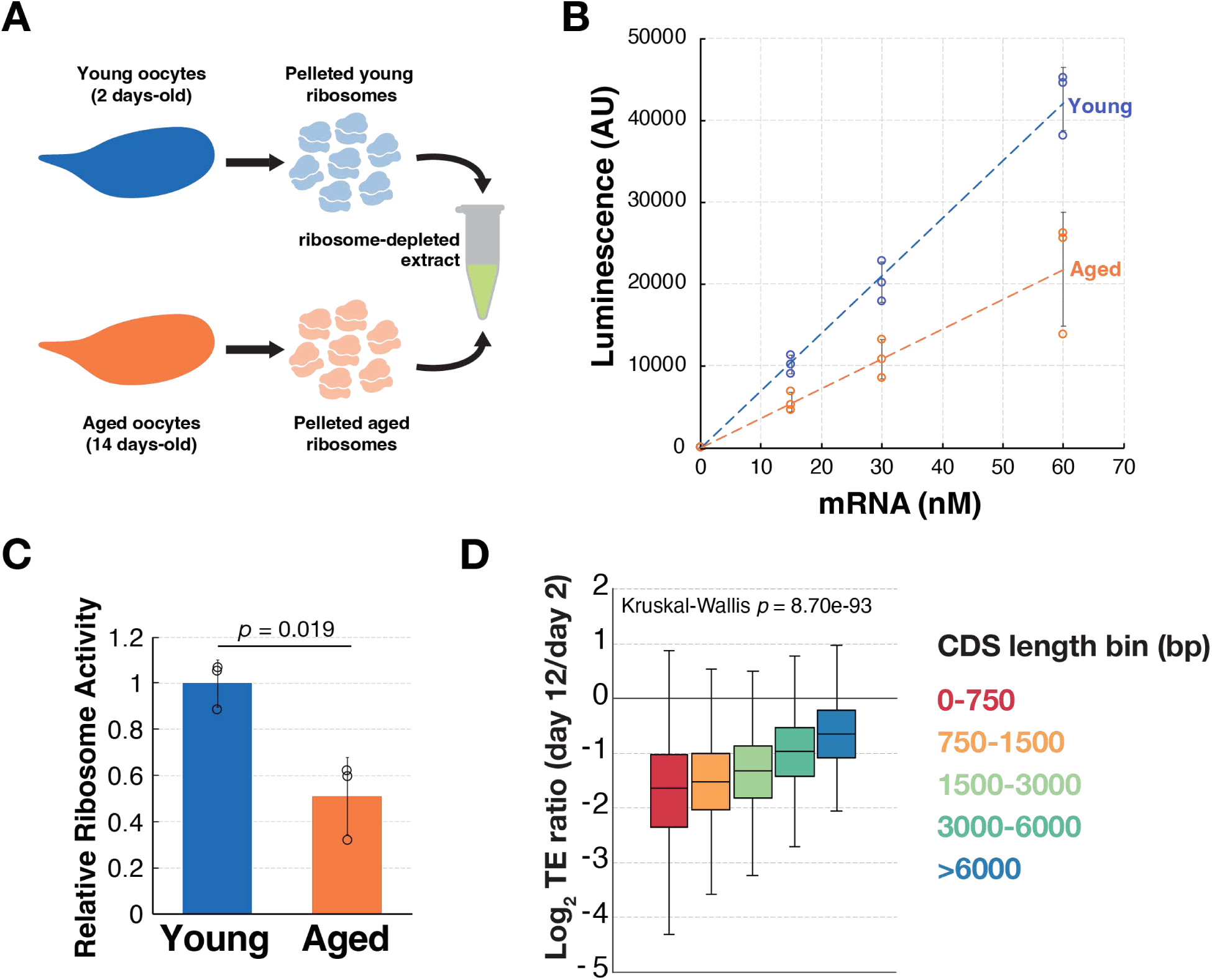
Ribosomes become less active in aged oocytes. **(A)** Schematic for *in vitro* translation assay using pelleted ribosomes from young (day 2) and old (day 14) oocytes. Ribosomes were added back into a ribosome-depleted extract to test activity in an identical environment. **(B)** Plot showing *in vitro* translation output of a luciferase cricket paralysis virus (CrPV) IRES reporter is reduced from ribosomes obtained from aged vs. young oocytes (n = 3 biological replicates per age). **(C)** Quantification showing day 14 ribosomes are ∼50% reduced in activity relative to day 2 (Welch’s unpaired *t*-test, p = 0.019). Bars represent group means; individual replicate slopes shown as points. **(D)** Plot showing translation efficiency decreases more for shorter transcripts during oocyte aging, similar to cells with reduced ribosomes (Khajuria et al., 2018).

We next asked whether aging oocytes exhibit an *in vivo* translational signature consistent with reduced functional ribosome availability. Reduced ribosome abundance can produce transcript-length-dependent effects on translation, with shorter transcripts disproportionately affected (Khajuria et al. 2018; Babaylova et al. 2025). We therefore re-analyzed previously published spike-in-normalized ribosome profiling data from aging oocytes (Greenblatt et al. 2019). Transcript length was strongly associated with the age-dependent change in translation efficiency, with shorter transcripts showing the greatest decline (Fig. 3D). Across coding regions, the reduction in ribosome footprint density was relatively uniform, with no systematic age-dependent shift in the positional distribution of ribosomes (Fig. S3). Together, the uniform reduction in ribosome occupancy across transcript bodies and the transcript-length dependence of translational decline are consistent with a reduction in the pool of functionally available ribosomes during oocyte aging.

### Both 40S and 60S ribosomal subunits lose activity during oocyte aging without detectable changes in rRNA structure or ribosomal protein composition

The decline in translation during *Drosophila* oocyte aging could reflect a reduction in ribosome abundance in addition to the loss of intrinsic ribosome activity described above. To quantify ribosome abundance, we measured rRNA levels using spike-in-normalized Bioanalyzer analysis to control for differences in RNA recovery between samples. Total rRNA abundance declined by approximately 15% between young and aged oocytes (Fig. 4A). Thus, *Drosophila* oocyte aging is accompanied by both a modest reduction in ribosome abundance and a 50% reduction in the intrinsic translational activity of the remaining ribosomes, explaining the substantial decrease in global translation rates previously observed (Greenblatt 2019).

**Figure 4.**
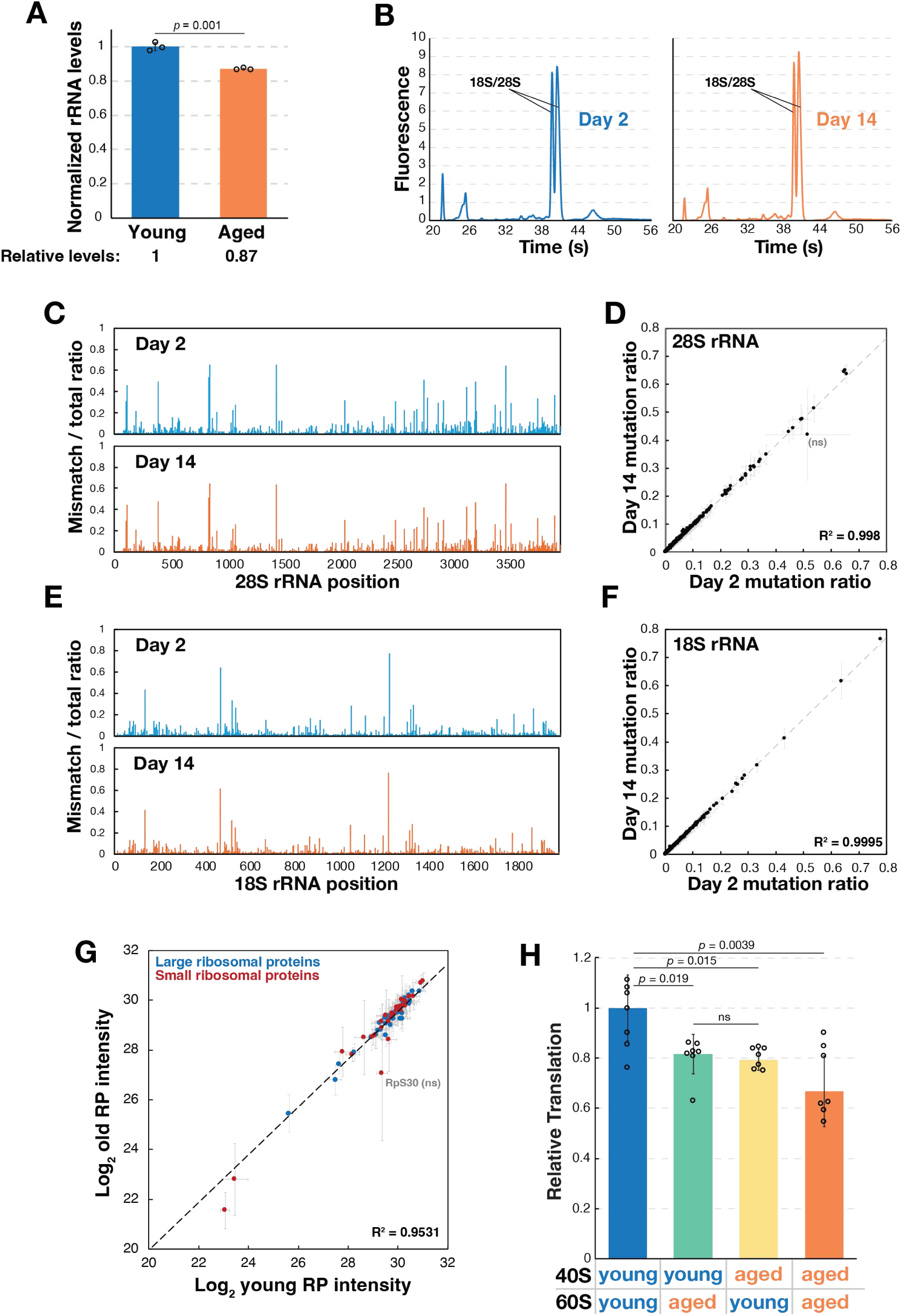
40S and 60S ribosomal subunits each show reduced activity during aging in the absence of observable changes to rRNA structure. **(A)** Quantification of bioanalyzer traces showing that rRNA abundance falls modestly (∼15%) with oocyte age. **(B)** Total RNA assessed by bioanalyzer shows no detectable cleavage of rRNA in day 2 or day 14 oocytes. **(C)** DMS-MaPseq of 28S rRNA in young (day 2, blue) and old (day 14, orange) shows no observable change in basepairing, indicative of stable rRNA folding. Mismatch/total ratio at each nucleotide position reflects DMS reactivity at adenines and cytosines. **(D)** Correlation of per-nucleotide DMS mutation ratios between young and old oocytes for 28S rRNA (R² = 0.998), indicating minimal structural change with age. **(E)** DMS mutational profiling of 18S rRNA in young and old oocytes, as in (C). **(F)** Correlation of per-nucleotide DMS mutation ratios between young and old oocytes for 18S rRNA (R² = 0.9995), as in (D). **(G)** Mass spectrometry of ribosomal protein abundance from pelleted ribosomes in young (day 2) versus old (day 14) oocytes from 4 replicates. Log_2_ intensity of individual ribosomal proteins in old relative to young oocytes, with linear regression (R² = 0.9531) shows that ribosomal protein stoichiometry is largely unchanged with age; RpS30, shows a non-significant (ns) deviation. **(H)** Subunit-swapping experiments between young and old preparations reveals that both 40S and 60S independently lose translational activity. Welch’s *t*-test was used for statistical analyses. N = 7 biological replicates, 2 technical replicates each.

We next investigated whether the reduced activity of aged ribosomes was associated with detectable changes in ribosome integrity or composition. Because mature oocytes must maintain stored RNA over prolonged periods, we first asked whether rRNA undergoes degradation during aging. Bioanalyzer analysis revealed no detectable fragmentation of rRNA in aged oocytes (Fig. 4B), arguing against extensive rRNA cleavage or ribosome degradation. In *Drosophila*, 28S rRNA a naturally occurring “hidden break” that generates two fragments similar in size to 18S rRNA (Natsidis et al. 2019); consequently, the absence of the conventional intact 28S peak in these electropherograms is characteristic of *Drosophila* rRNA rather than evidence of RNA degradation.

Because rRNA forms the catalytic and structural core of the ribosome, we considered whether age-dependent alterations in rRNA itself or in its interactions with ribosomal proteins might accompany the loss of ribosome activity. We therefore probed rRNA structure *in vivo* in young and aged oocytes using DMS-MaPseq (Zubradt et al. 2017). We optimized a DMS reactivity assay for 28S and 18S rRNAs (Fig. S4) and found that the reactivity profiles were highly similar between young and aged oocytes (Fig. 4C–F), with coefficients of determination of R² = 0.998 and R² = 0.9995, respectively. Thus, we detected no large-scale age-dependent changes in rRNA accessibility by DMS-MaPseq. Similarly, mass-spectrometric analysis of pelleted ribosomes revealed that ribosomal protein stoichiometry was largely preserved during aging, with no significant change in the relative abundance of individual ribosomal proteins with one exception, RpS15Ab, for which we could only identify one unique peptide due to its sequence similarity with the paralogous protein RpS15Aa (Fig. 4G, Fig. S5). Together, these results indicate that the substantial loss of ribosome activity cannot be explained by large-scale changes in rRNA structure or ribosomal protein composition.

Having found no gross structural or compositional change that could explain the decline in ribosome activity, we sought to determine whether the functional defect was preferentially associated with the small or large ribosomal subunit. We separated 40S and 60S subunits from young and aged oocytes by sucrose-gradient centrifugation and combined different combinations of young and aged subunits in the ribosome-depleted *in vitro* translation system described above. Ribosomes containing both aged 40S and 60S subunits exhibited ∼65% translational activity relative to those containing two young subunits (Fig. 4H; *p* = 0.0039, Welch’s *t*-test). The magnitude of this age-dependent reduction was somewhat smaller than that observed with unfractionated ribosomes (Fig. 3B,C), potentially reflecting differences introduced by the additional sucrose-gradient purification and subunit reconstitution steps. Replacing either the 40S or 60S subunit alone with an aged subunit produced an intermediate reduction in activity to ∼80% in both cases (*p* = 0.015 and *p* = 0.019 respectively, Welch’s *t*-test). These results indicate that age-dependent defects arise in both the 40S and 60S subunits and that each independently contributes to the overall decline in ribosome activity.

### Reduced ribosome content accelerates the development of meiotic and embryonic defects during oocyte aging

To determine why oocytes with reduced ribosome content lose developmental competence prematurely, we first asked whether they successfully complete meiosis and enter embryogenesis. In controls, most embryos derived from oocytes stored for 10 days progressed into embryogenesis, whereas after 17 days a large fraction arrested before embryonic development. In each case, the fraction of embryos that failed to enter embryogenesis closely paralleled the reduction in hatch rate, consistent with meiotic failure accounting for much of the age-dependent loss of viability. A similar premature arrest was observed in embryos derived from 10-day-old *RpL29^Df^*/+ oocytes, whereas a null-allele *cactus*^Mi^/+ phenocopied the hatching defect in the corresponding deficiency line but did not produce a comparable meiotic phenotype (Fig. 5A,B). Thus, reducing *RpL29* dosage selectively accelerates the appearance of a meiotic defect that normally emerges only after much longer periods of oocyte storage.

**Figure 5.**
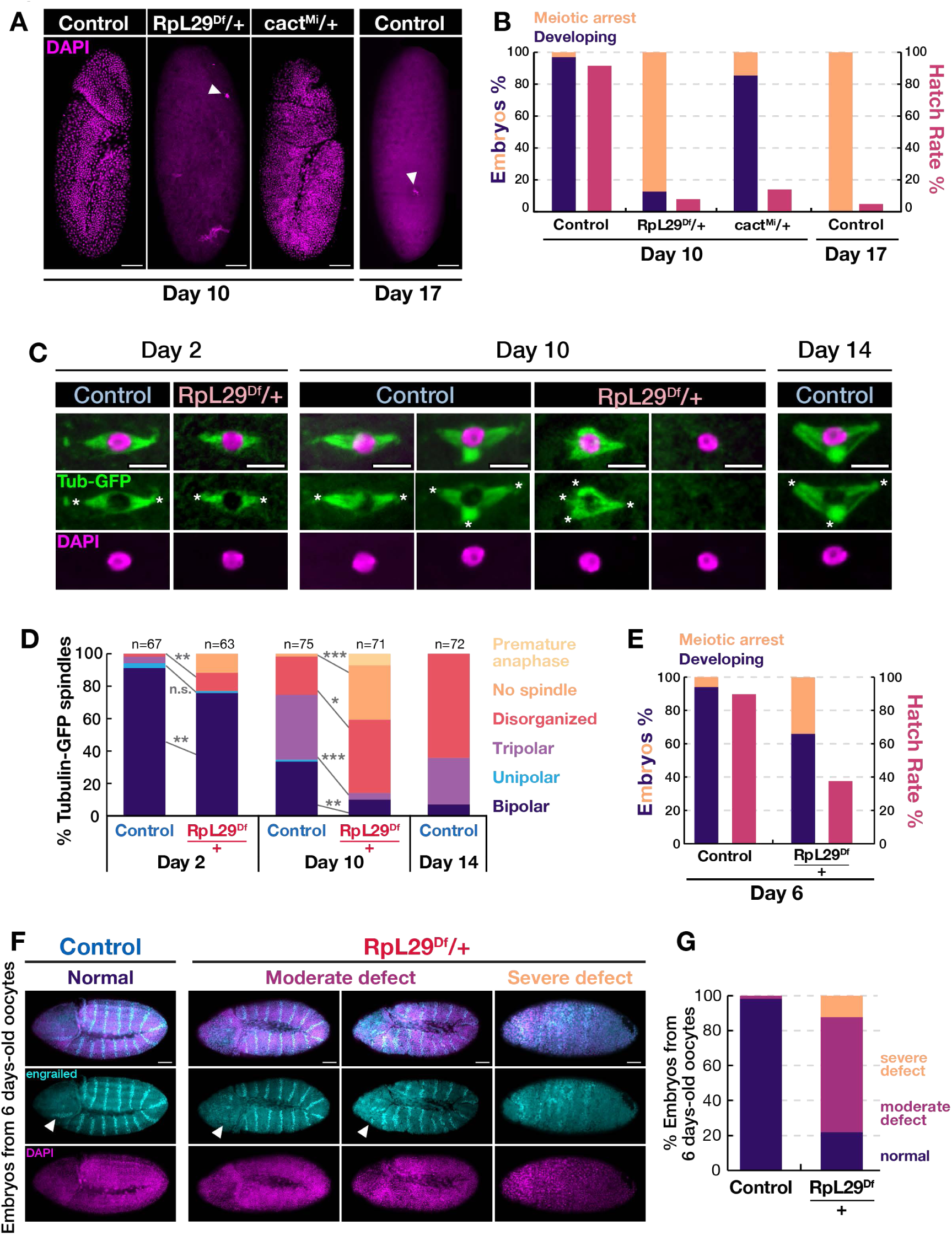
Oocytes haploinsufficient for *RpL29* prematurely develop meiotic and patterning defects with age. **(A)** Representative images of 4–8-hour embryos from stored control, *cact^Mi^*/+ (heterozygous for *cact* null-allele) *RpL29^Df^*/+ (heterozygous for the deficiency line containing *RpL29*) oocytes showing that reduced *RpL29* gene dosage impairs onset of embryonic development from aged oocytes, consistent with a meiotic defect. White arrow marks arrested polar bodies, scale bar is 50 µm. **(B)** Reduced *RpL29* but not *cactus* gene dosage substantially impairs meiotic function in aged oocytes. Quantification of the percentage of 4–8-hour embryos that progress through embryogenesis (“developing”) or arrest prior to embryogenesis (“meiotic arrest”), alongside their corresponding hatch rates for indicated genotypes. **(C)** Representative images of meiotic spindles labeled with Tubulin-GFP showing increased incidence of highly disorganized/absent spindles in *RpL29^Df^*/+ oocytes at various aging timepoints. Asterisk marks spindle poles, scale bar is 5 µm. **(D)** Quantification of meiotic spindle morphology. *RpL29^Df^*/+ oocytes show accelerated decline in spindle bipolarity and increased incidence of morphological defects with age. Welch’s unpaired *t*-test, *P < 0.05, **P < 0.005, ***P < 0.0005. **(E)** Quantification of embryos from 6 day-old *RpL29^Df^*/+ oocytes show that the percentage of developing embryos does not fully correlate with hatch rate at intermediate timepoints, suggesting additional defects may arise during embryogenesis that further reduce viability. **(F)** Immunostaining for the patterning protein engrailed in 4–8-hour embryos from 6-day-old control and *RpL29^Df^*/+ oocytes, showing clear bands in embryos from control oocytes, but exhibiting moderate and severe expression defects in embryos from *RpL29^Df^*/+ oocytes. White arrow marks the anterior-most engrailed band that is missing in *RpL29^Df^*/+ embryos. Scale bar is 50 µm. **(G)** Quantification of 4–8-hour embryos that progress past meiosis shows that the majority of embryos from 6-day-old *RpL29^Df^*/+ but not control oocytes have moderate to severe engrailed expression defects, indicative of impaired patterning.

We next tested whether reduced ribosome content in *RpL29^Df^*/+ oocytes accelerated deterioration of the metaphase-I spindle. Using Tubulin-GFP to visualize meiotic spindles, we found that most young *RpL29^Df^*/+ oocytes contained bipolar spindles, although the proportion was modestly (10-15%) reduced compared to controls. With increasing storage time, both genotypes showed a progressive loss of bipolar spindle morphology and an increase in spindle defects, but these changes occurred substantially earlier in *RpL29^Df^*/+ oocytes, as the fraction of spindles that retained bipolarity in 10-day old *RpL29^Df^*/+ oocytes was not significantly different from 14-day old controls (Fig. 5C,D, *p* = 0.2, Welch’s *t*-test). By 10 days of storage, *RpL29^Df^*/+ oocytes also showed a marked increase in oocytes lacking a detectable spindle, a phenotype not observed in control oocytes at the timepoints examined. Because translation is required for normal spindle assembly and maintenance (Susor et al. 2015), this accelerated loss of spindle integrity is consistent with reduced translational capacity compromising the machinery required to maintain the meiotic spindle during prolonged oocyte storage. Together, these results show that reducing ribosome content accelerates the age-dependent deterioration of meiotic spindle organization.

We next examined an intermediate storage time of 6 days, when *RpL29^Df^*/+ oocytes retained a hatch rate of approximately 38%. In control oocytes, the proportion of embryos that successfully entered embryogenesis (∼94%) closely matched the hatch rate (∼90%), indicating that embryos completing meiosis generally remained developmentally competent. In contrast, approximately 66% of embryos derived from *RpL29^Df^*/+ oocytes entered embryogenesis despite only ∼38% ultimately hatching (Fig. 5E), indicating that a substantial fraction failed after meiotic completion. We therefore examined embryonic patterning using Engrailed, which normally forms a stereotyped series of segmental stripes. Among embryos derived from 6-day-old *RpL29^Df^*/+ oocytes that progressed into embryogenesis, the majority exhibited moderate or severe abnormalities in Engrailed patterning, frequently including loss of the anterior-most stripe (Fig. 5F,G). Thus, reduced ribosome content produces defects at multiple stages: meiotic failure becomes increasingly prominent with age, while at intermediate timepoints embryos that successfully complete meiosis can nevertheless exhibit substantial defects during subsequent development.

### Reduced ribosome content accelerates age-dependent defects in spindle poles and kinetochores

To define which features of spindle organization are affected by reduced ribosome content, we examined spindle poles and kinetochores using Klp10A-GFP and Cenp-C-GFP transgenes, respectively. We first assessed spindle pole organization. Young *RpL29^Df^*/+ oocytes showed a modest reduction in the proportion of spindles with two clearly defined Klp10A-positive poles compared with controls (Fig. 6A,B), despite retaining near-normal developmental competence (Fig. S6A). With increasing age, however, *RpL29^Df^*/+ oocytes showed an accelerated loss of normal bipolar Klp10A localization and a corresponding increase in spindles with a single pole or multiple/poorly defined poles (Fig. 6A,B), as the fraction of double-Klp10A-positive spindles was not significantly different between 10-day old *RpL29^Df^*/+ oocytes and 16 day-old control oocytes (*p* = 0.2, Welch’s *t*-test). Thus, defects in spindle pole organization emerge prematurely in oocytes with reduced ribosome content and accompany the broader age-dependent deterioration of spindle morphology.

**Figure 6.**
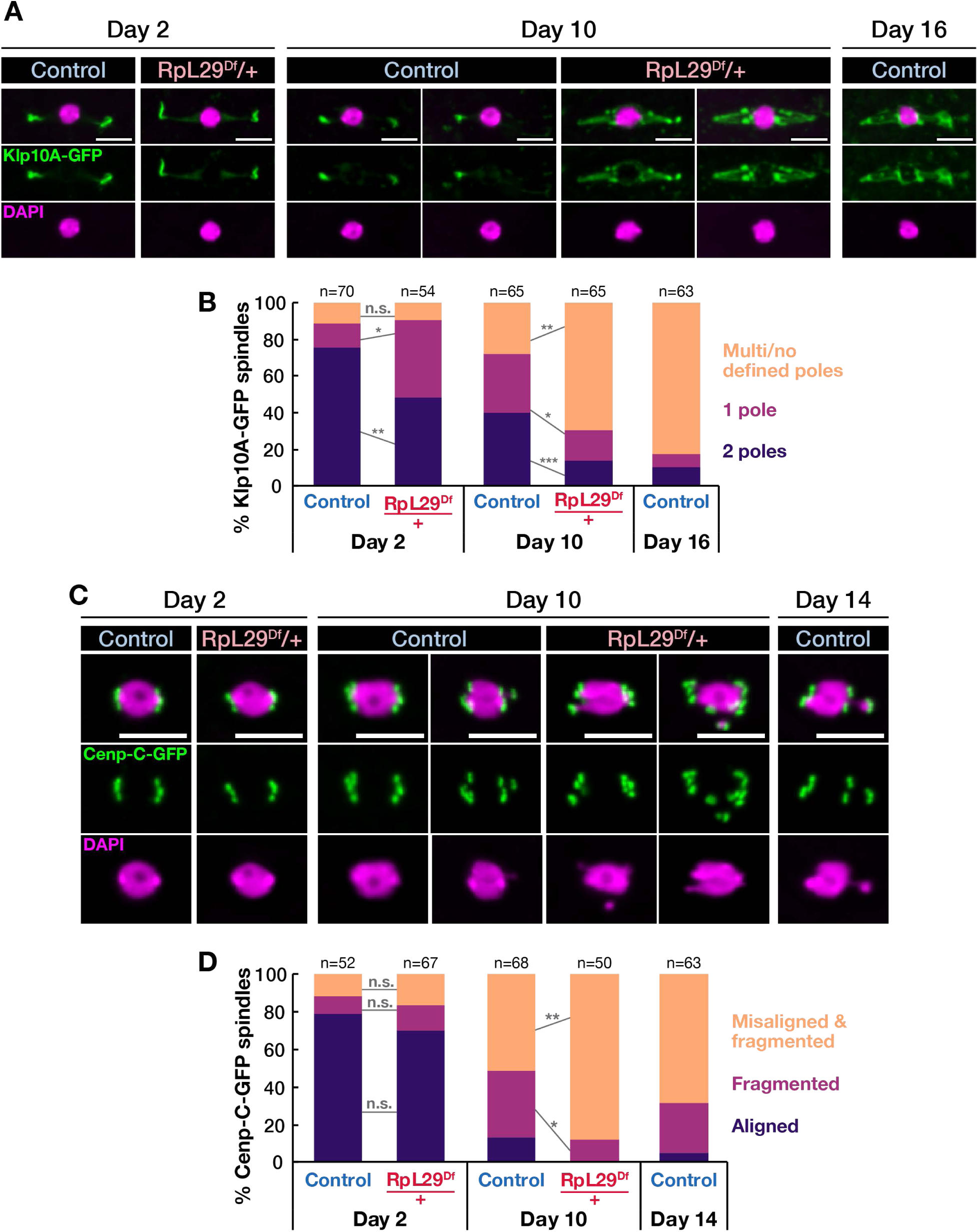
Oocytes haploinsufficient for *RpL29* prematurely develop spindle pole and kinetochore defects with age. **(A)** Representative images showing defective Klp10A-GFP labeling in meiotic spindle poles from *RpL29^Df^*/+ and control oocytes after 2, 10, and 16 days of aging. **(B)** Quantification of Klp10A-GFP spindle pole morphology with age, scored as “2 poles” (two clearly defined poles), “1 pole” (a single clearly defined pole), or “multi/no defined poles” (multiple poles or no clearly defined poles). *RpL29^Df^*/+ oocytes show accelerated decline in normal spindle pole morphology with age, with a rise in multi/no defined poles category. **(C)** Representative images of Cenp-C-GFP, showing impaired kinetochore distribution during aging in *RpL29^Df^*/+ oocytes after 2, 10, and 14 days. **(D)** Quantification of Cenp-C-GFP kinetochore morphology with age, scored as “aligned” (3–4 clearly defined kinetochores), “fragmented” (fragmented Cenp-C signal, i.e., more than 4 kinetochore foci, with normally aligned DNA), or “misaligned & fragmented” (fragmented Cenp-C signal together with misaligned DNA that fails to condense into a single round mass). *RpL29^Df^*/+ oocytes show accelerated decline in normal kinetochore morphology with age, with a rise in misaligned & fragmented category. Welch’s unpaired *t*-test, *P < 0.05, **P < 0.005, ***P < 0.0005. Scale bar is 5 µm.

We next assessed kinetochore organization using Cenp-C-GFP. Kinetochore morphology was similar between young control and *RpL29^Df^*/+ oocytes, indicating that reduced ribosome content does not produce a substantial baseline defect in kinetochore organization. The GFP tag itself had a destabilizing effect, as Cenp-C-GFP animals exhibited premature loss of oocyte function (Fig. S6B). With age, however, *RpL29^Df^*/+ oocytes showed an accelerated loss of normal Cenp-C-GFP organization accompanied by increased kinetochore fragmentation and the appearance of fragmented kinetochores associated with chromosome misalignment (Fig. 6C,D). Thus, reduced ribosome content can accelerate age-dependent defects in both spindle pole and kinetochore organization, indicating that multiple features of meiotic spindle architecture are sensitive to diminished translational capacity.

### Protein synthesis declines in aged metaphase I mouse oocytes

Previous studies have also reported reduced protein synthesis in aged mouse oocytes either before meiotic resumption at the so-called germinal-vesicle stage (meiotic prophase I arrest) or at metaphase II, after completion of meiosis I (Zhang et al. 2026; Takahashi et al. 2023; Huang et al. 2023). Because meiosis I is when the meiotic spindle is assembled and the majority of age-related chromosome segregation errors arise (Sakakibara et al. 2015), we compared protein synthesis in metaphase I oocytes. Young and aged mouse oocytes were matured *in vitro* to metaphase I and incubated with the methionine analog L-homopropargylglycine (HPG), followed by fixation and click-chemistry-based fluorescent labeling of newly synthesized proteins (Fig. 7A). Cycloheximide-treated oocytes were analyzed in parallel to define the translation-dependent component of the HPG signal. Aged metaphase I oocytes exhibited markedly reduced HPG incorporation relative to young oocytes, with signal approaching that observed following cycloheximide treatment (Fig. 7B,C). Cycloheximide treatment severely impaired spindle formation, whereas aged oocytes retained spindles with volumes comparable to those of young oocytes (Fig. 7D,E). These observations indicate that aged metaphase I mouse oocytes retain low-level residual translational activity that is much reduced as compared to oocytes from young animals.

**Figure 7.**
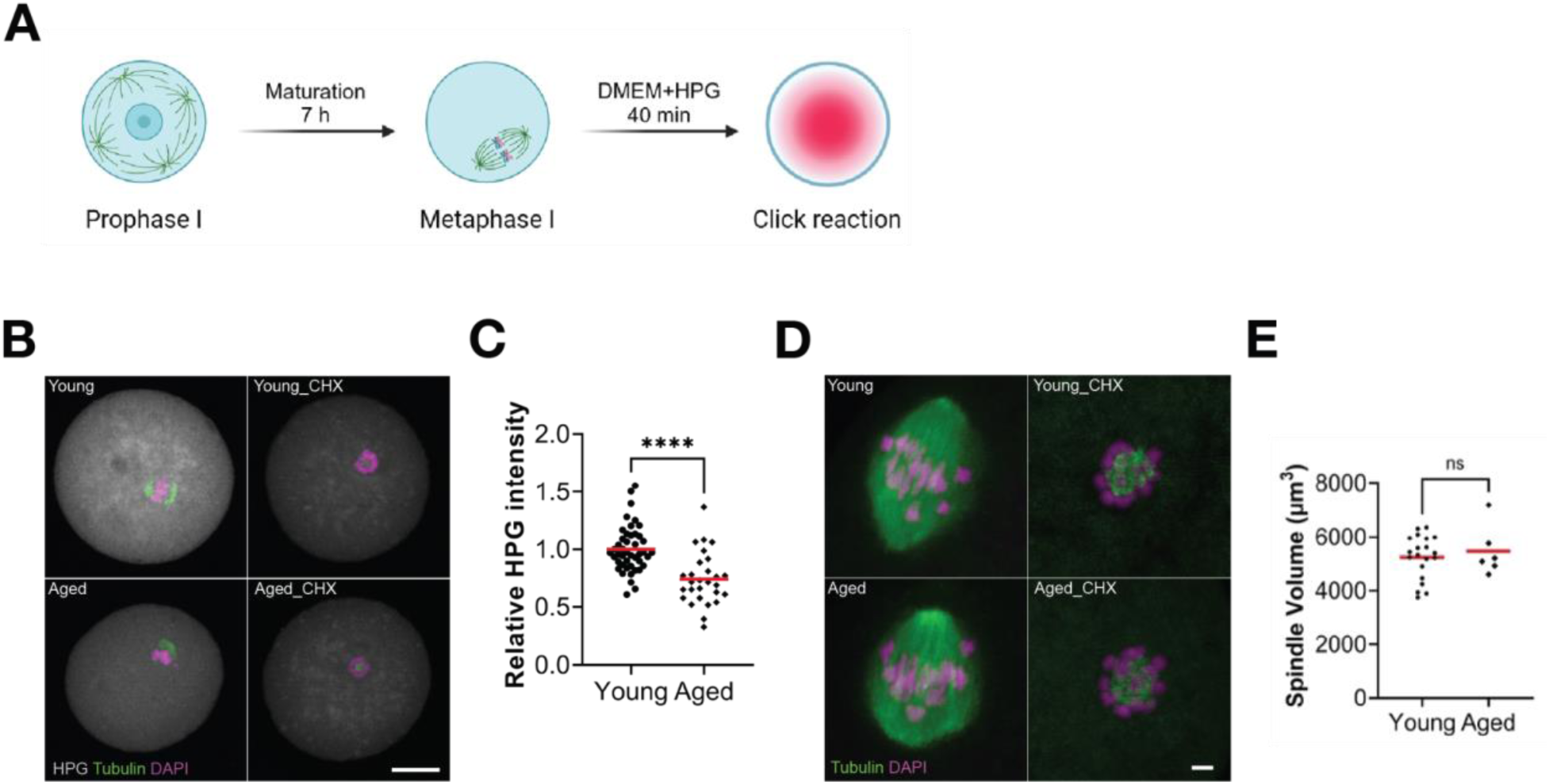
Protein synthesis declines in aged metaphase I mouse oocytes. **(A)** Schematic of HPG (L-homopropargylglycine) translation assay. **(B)** Representative confocal images of metaphase I (Met I) oocytes assessed using HPG click-reaction in the young and aged groups. Scale bars: 20 μm. **(C)** Quantitation of HPG intensity, normalized to the mean of young oocytes. The experiment was performed twice. Data points represent individual oocytes. Young, n = 45 and aged, n = 28. Red horizontal line indicates the mean. Differences between the two groups were analyzed using a two-tailed unpaired Student’s *t*-test. \*\*\*\**P* < 0.0001. **(D)** Representative confocal images of Met I spindle in the indicated groups showing that translation is required for spindle formation. Scale bars: 5 μm. CHX treatment impaired spindle formation. **(E)** Spindle volume quantification in young and aged oocytes. Differences were analyzed using Student’s *t*-test. Each dot represents an individual oocyte. Red horizontal line indicates the mean.

## DISCUSSION

Our genetic and biochemical analyses suggest that ribosome dysfunction is a major contributor to age-dependent meiotic spindle instability in *Drosophila* oocytes. This vulnerability may extend to mammals, where we and others observe a striking decline in protein synthesis during meiotic maturation. Although mammalian oocytes do not undergo the prolonged metaphase I arrest characteristic of *Drosophila*, both systems rely on a highly conserved developmental program in which ribosomes are stockpiled during oocyte growth for later use. This reliance on a long-lived, pre-existing ribosome pool may create an inherent vulnerability to aging. We find that ribosome activity declines substantially with age, with both the 40S and 60S subunits independently contributing to this loss of function. Yet this functional decline occurs without detectable large-scale changes in rRNA structure or ribosomal protein stoichiometry, suggesting that more subtle alterations, such as chemical modification or accumulated molecular damage, underlie the loss of activity. One possibility is oxidative damage. Ribosomes are established targets of oxidation (Shcherbik and Pestov 2019), and redox imbalance itself disrupts meiosis and early embryonic divisions (Petrova et al. 2018; Perkins et al. 2016). The prolonged lifetime of oocyte ribosomes may therefore increase their susceptibility to the cumulative effects of such cellular stresses.

Previous studies have identified age-related defects in spindle assembly checkpoint activity, kinetochore function, and chromosome cohesion, but whether these changes arise independently or reflect a common upstream decline remains unclear. One potential unifying mechanism is impaired translation. In mammalian oocytes, translation is spatially activated around chromosomes following nuclear-envelope breakdown through the mTOR–eIF4F pathway and is required for normal spindle assembly, chromosome alignment, and segregation (Susor et al. 2015; Severance and Latham 2017; Romasko et al. 2013). Maternal aging also disrupts the CPEB1-dependent translational program: aged mouse oocytes show reduced CPEB1 expression and altered translation of its targets, while partial CPEB1 loss in young oocytes recapitulates several age-associated translational and meiotic phenotypes (Takahashi et al. 2023). Together, these studies indicate that precisely regulated translation is critical for meiotic progression and becomes compromised with age. Consistent with this model, multiple spindle defects co-occur in aged wild-type *Drosophila* oocytes, and experimentally reducing ribosome abundance accelerates the appearance of each phenotype. Thus, declining translational capacity may represent a common upstream defect that broadly compromises chromosome segregation machinery during oocyte aging.

Reduced translational capacity may also compromise developmental competence beyond meiosis. At intermediate ages, many *RpL29^Df^*/+ oocytes successfully completed meiosis, yet the resulting embryos subsequently developed severe patterning defects, indicating that reduced ribosome abundance can impair maternal functions required during early embryogenesis. This broader effect is consistent with emerging evidence that maternal age produces embryo-intrinsic defects that extend beyond aneuploidy. Recently, Cavanaugh et al. showed that embryos from aged mice exhibit increased trophectoderm contractility, tissue surface tension, and viscosity, which impair embryo spreading and implantation; related mechanical changes were also observed in human embryos from older women (Cavanaugh et al. 2026). Although the molecular basis of these mechanical defects remains unclear, our findings raise the possibility that age-dependent deterioration of maternal translational capacity could contribute not only to meiotic failure but also to defects in the cellular programs that support subsequent embryonic development.

These findings also have implications for interventions targeting nutrient-sensing pathways to extend reproductive lifespan. mTOR inhibition has been proposed as a strategy to preserve ovarian reserve by limiting primordial follicle activation. However, mTOR signaling also promotes translation during meiotic resumption and supports spindle assembly through the mTOR–eIF4F axis (Susor et al. 2015). Our finding that reduced ribosome abundance accelerates *Drosophila* oocyte dysfunction raises the possibility that prolonged or poorly timed mTOR inhibition could impose a tradeoff between preserving the dormant follicle pool and maintaining the translational capacity required for oocyte maturation. Interventions attempting to preserve follicle number may not necessarily preserve overall ovarian function (Bonus et al. 2024), emphasizing the importance of timing and duration when targeting mTOR in the aging ovary.

Our findings also raise the possibility that some features of oocyte aging may be reversible through restoration of limiting cellular components. Recent experiments in aged mouse oocytes showed that exposure to soluble nuclear factors from young oocytes substantially reduced age-associated DNA damage, improved chromosome segregation, and restored developmental competence, demonstrating that some defects associated with oocyte aging can be ameliorated by replenishing components of the young oocyte (Dudko et al. 2026). Similarly, cytoplasmic transfer from young donor oocytes has been reported to improve fertilization and early embryonic development in human oocytes of poor quality, although the responsible components remain unknown (Sobek et al. 2020). Our results identify translational capacity as another potentially limiting feature of the aged oocyte and raise the possibility that preserving or restoring functional ribosomes could improve meiotic and developmental competence. More broadly, these findings support a model in which reproductive aging reflects not only irreversible damage to chromosomal complexes, but also the progressive deterioration of long-lived gene expression machinery that may, in principle, be preserved or restored.

## METHODS

### Transgenic animals and husbandry

Flies were maintained on cornmeal–yeast–dextrose medium at 25 °C. The medium contained 70.9 g D-glucose (dextrose), 48.5 g cornmeal, 30.3 g active dry yeast, 20.5 g sucrose, 4.55 g agar, 0.5 g CaCl₂·2H₂O, and 0.5 g MgSO₄·7H₂O, cooked in approximately 0.47 L water and brought to final volume. After cooling, a propionic/phosphoric acid preservative mixture was added to final concentrations of 0.49% (v/v) propionic acid and 0.049% (v/v) phosphoric acid.

The RpL29 rescue construct was generated in the UASz1.0 backbone. DNA fragments encoding the full-length coding sequences of RpL29 were synthesized and inserted into the BamHI and XbaI sites of UASz1.0 (GenScript). Constructs were injected into the attP40 landing-site lines (Genome Prolab Inc.) for φC31 integrase–mediated integration, and transformants were identified by screening for the mini-white (w⁺) eye-color marker.

### Oocyte isolation

Oocytes were aged as in Greenblatt et al. (2019). *Drosophila* ovaries were dissected in Grace’s Insect Media (Sigma-Aldrich) from flies fed for two days with wet yeast paste and oocyte storage was induced by placing animals onto molasses media (44 g agar, 180 mL molasses, 37 mL in 5% Tegosept, 1112 mL water) for the desired number of days. Stage 14 oocytes were isolated with 1 KU collagenase (Worthington, LS005273) in Grace’s + 0.1% Triton-X for 15 min at room temperature with gentle agitation and washed four times with Grace’s + 0.1% Triton-X. Oocytes were rinsed on a 70 µm strainer and rinsed three times with Grace’s to remove debris and eluted from the filter and washed four times with DM buffer (24 mM HEPES pH 7.5, 80 mM KOAc, 1 mM Mg(OAc)_2_).

### Genetic Screening and Hatch Rate Assay

A full list of experimental genotypes and fly stock sources is provided in Supplementary Table S2. Stocks used in the deficiency screen were from the Bloomington 2R and 2L kit and outcrossed with Oregon R wild-type strain, specific lines listed in Supplementary Table S3. Hatch rate assay was performed at 25 °C according to (Greenblatt et al. 2019). Number of days-old listed is the number of days spent on agar-sugar food (44 g agar, 180 mL molasses, 37 mL of 5% Tegosept, 1112 mL water). Flies for young oocytes were not put on molasses while aged were on molasses vials for 10 days.

### Egg laying assay

Three males (3–7 days old and previously isolated from females for at least 2 days) and three newly eclosed virgin females were placed in a bottle with a scored molasses plate topped with wet yeast paste and allowed to lay eggs for 24 hours at 25°C before the plate was changed. The plate was then replaced daily, and eggs were counted and normalized to the number of females present.

### rRNA quantification

Individual oocytes were counted and lysed in lysis buffer (15 mM Tris-HCl pH 7.5, 300 mM NaCl, 6 mM MgCl₂, 1% Triton X-100, 1 mg/mL heparin, 1 mM DTT) at 1 µL buffer per oocyte, using a dounce homogenizer. A 5 µL aliquot of extract was spiked with 250 ng of CrPV IGR IRES luciferase RNA (3597 nt) prior to total RNA extraction with TRIzol. Extracted RNA was analyzed on an Agilent Bioanalyzer, and rRNA peak area was normalized to spike-in peak area to control for differences in extraction efficiency across samples. 18S and 28S rRNA were quantified from the 2 rRNA peaks on the electropherogram traces. Because *Drosophila* 28S rRNA is cleaved into two fragments during rRNA processing, it co-migrates with 18S rRNA, appearing as two closely spaced peaks. The first peak was used to quantify 28S rRNA, and 18S rRNA levels were calculated by subtracting the first peak’s signal from that of the second peak.

### SP3 and DIA-Mass Spectrometry

5 µg of crude ribosomal extract was digested using the single-pot sample preparation protocol (Hughes et al. 2019) with SeraMag^TM^carboxylate-modified beads SpeedBeads [E7] and [E3] (45152105050250 and 65152105050250, Cytiva, Wilmington, DE, USA) and sequencing-grade modified trypsin (V51135, Promega, Madison, WI, USA). Briefly, samples were first treated with 3 mM TCEP (tris(2-carboxyethyl) phosphine) (25°C for 20 minutes) then with 55 mM 2-chloroacetamide (25°C for 30 minutes). Magnetic beads were added to the samples at 1:10 (protein:bead) ratio in a 50:50 (v/v) mixture with 100% ethanol (24°C for 10 minutes), then washed thrice with 80% ethanol before incubation with 200 ng of trypsin in 50 mM HEPES (pH = 8.0) at 37°C for 14 hours. 10%TFA was added to supernatants prior to desalting with the stop and go extraction (StAGE) (Rappsilber et al. 2003) tipping using an AssayMAP Bravo (Agilent, Santa Clara, CA, USA) liquid handler and AssayMAP 5 µL C18 cartridges (5190–6532, Agilent, Santa Clara, CA, USA). The peptides were eluted with 0.1% TFA, 40% ACN before drying and then reconstituted in 0.1% formic acid (FA) and 0.5% ACN before mass spectrometry.

The peptides were analyzed with an Orbitrap Exploris 480 mass spectrometer (Orbitrap Exploris^TM^ 480, Thermo Fisher Scientific) coupled to an Easy-nLC 1200 (Thermo Fisher Scientific) with Aurora Series analytical column (Ion Opticks) and Nanospray Flex^TM^ ion source, and operated in a data-independent (DIA) mode. Full scans MS resolution was set to 60,000 with 300% normalized automatic gain control (AGC) target, 50% RF lens, 25 ms maximum injection time and scan range from m/z 380 Th to m/z 985 Th. DIA fragment spectra were collected at a resolution of 15,000, normalized AGC target of 2000%, maximum injection time of 40 ms, scan range from m/z 145 Th – m/z 1450 Th. Isolation windows of m/z 10 Th were used with an overlap of m/z 1 Th. Normalized collision energy was set to 28%. The RAW files were searched with DIA-NN 2.0.2 and a *D. melanogaster* FASTA database from Uniprot with Q.Value < 0.01, PG.Q.Value < 0.05, Lib.Q.Value < 0.01 and Lib.PG.Q.Value < 0.01 and PG.MaxLFQ quantification. Missing values were imputed using the bottom 1% of intensity values. This was done in R, using the imputeLCMD package.

Protein intensity values were filtered to remove highly abundant yolk proteins (Vit1, Vit2, Vit3) and a trypsin (CON_TRYP_PIG) prior to analysis. Proteins with more than one imputed value across replicates within any condition were excluded from downstream analysis. Remaining intensity values were log2-transformed and normalized by subtracting each sample’s median intensity and adding the grand median across all samples, correcting for between-sample loading differences. Differential protein abundance was assessed between young and old oocytes, and young and RpL29^df^/+ oocytes, using Welch’s two-sample t-test for each protein, and p-values were adjusted for multiple testing using the Benjamini–Hochberg false discovery rate (FDR) procedure. Proteins were considered significantly differentially abundant at an FDR-adjusted p-value < 0.05 and an absolute log2 fold change > 1.

### Ribosome pelleting

Oocytes were weighed and dounce homogenized in Buffer D2 (20 mM Tris-HCl pH 7.5, 0.2 mM EDTA, 10 mM KCl, 10 mM MgCl₂, 6.8% sucrose, 1 mM DTT, 1 µL/mL SUPERase•In) at 4 µL buffer per 1 mg oocyte weight, using 15-20 strokes in a dounce homogenizer. The homogenate was cleared by centrifugation at 16,000 × g for 10 min at 4°C. The resulting supernatant was centrifuged at 140,000 × g for 1 hr at 4°C in a TLA-110 rotor to pellet ribosomes. The pellet was overlaid with 100 µL of Buffer D2, which was then removed without disturbing the pellet, to remove residual supernatant. The washed pellet was gently resuspended in Buffer D2 at 2 µL per 1 mg of starting oocyte weight.

### Ribosomal 40S and 60S subunit sucrose gradient purification

Ovaries were hand-dissected from 150 flies and stage 14 oocytes isolated as described above. Oocytes were dounce homogenized in 0.5 mL lysis buffer (15 mM Tris-HCl pH 7.5, 300 mM NaCl, 6 mM MgCl₂, 1% Triton X-100, 1 mg/mL heparin, 1 mM DTT) and clarified by centrifugation at 23,000 × g for 30 min at 4°C. The supernatant was layered over 1 mL of a sucrose cushion (34% sucrose, 0.5 M KCl, 2 mM DTT, 1 mM EDTA, 20 mM Tris-HCl, 6 mM Mg(OAc)₂, 1:1000 SUPERase•In) and centrifuged at 80,000 rpm for 1 hr 40 min at 4°C in a TLA-110 rotor to pellet ribosomes. The pellet was resuspended in 500 µL Buffer B (20 mM Tris-HCl pH 7.5, 6 mM Mg(OAc)₂, 150 mM KCl, 6.8% sucrose, 2 mM DTT).

To release bound endogenous mRNA, ribosomes were treated with 2.3 mM puromycin and sequentially incubated at 4°C for 10 min, 37°C for 10 min, and on ice for 10 min. KCl was then added to a final concentration of 350 mM, and the sample was incubated on ice for 10 min to dissociate 80S ribosomes into 40S and 60S subunits. Dissociated subunits were resolved on a 20–40% (w/w) sucrose gradient in Separation buffer (20 mM Tris-HCl pH 7.5, 0.5 M KCl, 6 mM MgCl₂, 2 mM DTT) by centrifugation at 22,000 rpm for 17 hr at 4°C in an SW-40 rotor (Beckman Coulter). Gradients were fractionated with continuous monitoring of absorbance at 260 nm to identify 40S and 60S peaks. Peak fractions were pooled and concentrated into Buffer D (20 mM Tris-HCl pH 7.5, 0.2 mM EDTA, 10 mM KCl, 1 mM MgCl₂, 6.8% sucrose) using Amicon Ultra 100 kDa MWCO concentrators (Millipore, UFC910008). Concentrations were determined spectrophotometrically using the conversions 1 A₂₆₀ = 50 nM for 40S and 1 A₂₆₀ = 25 nM for 60S.

### Preparation of ribosome-depleted extracts for *in vitro* translation

Oocytes were dounce homogenized in DM buffer supplemented with 2.5% RNase inhibitor (Cellscript, C-SRI6310K) at 4 µL buffer per 1 mg oocyte weight. The homogenate was centrifuged at 16,000 × g for 10 min at 4°C to remove debris. The resulting supernatant was centrifuged at 140,000 × g for 1 hr at 4°C in a TLA-110 rotor to remove ribosomes. Aliquots of the ribosome-depleted supernatant were stored at −80°C until use.

### Reporter RNA production using *in vitro* transcription

The luciferase reporter used in this study encodes an upstream Renilla luciferase reporter and a downstream firefly luciferase reporter which is translated by the presence of a CrPV IRES. To prevent translation of firefly luciferase by read-through or re-initiation, an inactivated encephalomyocarditis virus (EMCV) IRES is inserted upstream of the CrPV IRES. DNA was linearized with BamH1 restriction enzyme and *in vitro* transcribed with T7 RNA polymerase (New England Biolabs, E2040S) and post-transcriptionally 5’ capped (Cellscript, C-SCCE0625) and polyA-tailed (CellScript, C-PAP5104H). RNA integrity was monitored by TapeStation.

### *In vitro* translation assays

The in vitro translation protocol was adapted from methods developed for whole ovary extracts (Lie and Macdonald 2000). Each 10 µL reaction contained 24 mM HEPES-KOH pH 7.5, 1.5 mM Mg(OAc)₂, 80 mM KOAc, 0.75 mM ATP, 0.1 mM GTP, 0.2 mM spermidine, 16.8 mM creatine phosphate, 1.2 mM DTT, 60 µM complete amino acid mixture, 80 ng/µL creatine phosphokinase, 2.5% RNase inhibitor, variable amounts of in vitro-transcribed CrPV IGR IRES RNA, and 4 µL of ribosome-depleted extract. For pelleted ribosome experiments, 15 A₂₆₀ units of ribosomes were added per reaction; for purified subunit experiments, 30 nM each of purified 40S and 60S subunits were added instead. Reactions were carried out in 384-well plates (Corning) at 25°C for 90 min. Luciferase reagent (Promega) was then added (10 µL per reaction), and luminescence was measured after 2 min using a luminometer.

### Analysis of translation efficiency by transcript length

Ribosome profiling and mRNA-seq TPM values, along with pre-calculated translation efficiency (TE) values, were obtained from Greenblatt et al. (2019). Transcripts with a mean mRNA-seq TPM < 1 in either condition (day 2 or day 12) or a mean ribosome profiling TPM < 1 in either condition were excluded. CDS lengths for all *Drosophila melanogaster* genes were retrieved from Ensembl via the pybiomart Python package (v.0.2.0). Transcripts without a matching CDS length in Ensembl were excluded, resulting in 5,312 transcripts for final analysis. The change in TE was expressed as log₂(TE day12 / TE day2). Transcripts were grouped into five size classes based on CDS length: <750, 750–1500, 1500–3000, 3000–6000, and >6000 bp. Differences in log₂ TE ratio across size classes were assessed using the Kruskal-Wallis test, followed by pairwise Mann-Whitney U tests with Bonferroni correction.

### DMS-MaPSeq rRNA structural profiling

*In vivo* DMS modification of rRNA from stage 14 oocytes was adapted from Zubradt et al. 2017. Fifteen flies were dissected and oocytes isolated as described above, then suspended in 250 µL Grace’s media plus 250 µL DMS for 5 min at 25°C with shaking at 1000 rpm. Mock-treated samples received 250 µL Grace’s media in place of DMS. The reaction was quenched with 1 mL of 30% β-mercaptoethanol (BME), transferred to a 70 µm strainer, and washed three times with 30% BME and twice with sterile water. Oocytes were then collected and resuspended in 1 mL TRIzol for total RNA extraction.

DMS-MaPSeq libraries were prepared using the NEBNext UltraExpress RNA Library Prep Kit (New England Biolabs) according to the manufacturer’s protocol, with modifications adapted from Allan et al. 2024. Briefly, 100 ng of DMS-treated or mock-treated RNA was fragmented in 5 µL of 1X Fragmentation Buffer for 13 min. During first-strand synthesis, 1 µL of Induro Reverse Transcriptase (New England Biolabs) was substituted for the NEBNext UltraExpress First Strand Enzyme Mix, with incubated at 25°C for 10 min, 57°C for 30 min, and 95°C for 2 min. AMPure XP beads (Beckman Coulter) were used for all clean-up steps. Libraries were sequenced using 150-bp paired-end reads. Sequencing data were analyzed with SEISMIC-RNA to compute per-nucleotide mutation rates between DMS-treated and mock-treated samples (Allan et al. 2024).

### Embryo collection and Immunostaining

OregonR males (3–7 days old, isolated from females for at least 2 days) and aged females were mated for 4 hours at 25°C on scored agar–sugar plates (44 g agar, 180 mL molasses, 37 mL of 5% Tegosept, 1112 mL water), then incubated for an additional 4 hours to yield 4–8 hour embryos, which were collected into 1.5 mL tubes. Embryos were dechorionated for 2–3 min in 50% bleach, then washed in PBST (0.2% BSA, 0.1% Triton X-100 in 1× PBS).

Embryos were fixed for 25 min in a 1:1 mixture of fixative (1× PBS, 8% formaldehyde) and heptane on a nutator. The lower fixative layer was removed and replaced with an equal volume of methanol. Embryos were devitellinized by vortexing for up to 2 min, then the upper heptane layer was removed, and shook by hand for an additional 2 min. Embryos were washed three times in methanol and stored at −20°C until use.

Embryos were rehydrated in 50% methanol in PBST and washed three times in PBST. Embryos were blocked for 1 hour in PBST, then incubated overnight at 4°C with primary antibody (mouse anti-Engrailed, DSHB, 1:25, cat# ab12454) diluted in PBST. Embryos were washed three times in PBST for at least 20 min each and incubated overnight at 4°C with secondary antibody (CF568-conjugated goat anti-mouse, Biotium, 1:1000, cat # 20100) diluted in PBST. Embryos were washed three times in PBST for at least 20 min each, with DAPI added to the last wash, and mounted in 50% glycerol in 1× PBS.

### Spindle visualization

Ovaries were hand-dissected in room temperature Grace’s media. Ovaries were transferred into 1.5 mL tubes and fixed in 4% PFA in PBST for 25 min with rotation. Ovaries were washed twice for 20 min each in PBST, then incubated with DAPI (1:20,000) in PBST for 20 min with rotation. Ovaries were mounted in 50% glycerol in 1× PBS and sealed with nail polish. Images were acquired on an Olympus IXplore SR inverted spinning-disc confocal microscope using a 60× oil-immersion objective.

Spindle morphology was classified based on Tubulin-GFP, Klp10A-GFP, and Cenp-C-GFP localization patterns. Tubulin-GFP spindles were classified as bipolar (tubulin signal converging to two clearly defined poles), unipolar (convergence to a single pole), tripolar (convergence to three clearly defined poles), or disorganized (three or more poles forming without clear definition, tubulin signal present without defined poles, or tubulin association with an additional pole-like focus at the DNA). Spindles lacking detectable tubulin-GFP signal near the DNA were classified as no spindle, and those showing clear separation of DNA masses were classified as premature anaphase. Klp10A-GFP localization was classified as two poles (two clearly defined foci at opposite ends of the spindle), one pole (a single clearly defined focus), or multiple/no defined poles (mislocalization of Klp10A-GFP signal along the spindle, or absence of clearly defined foci). Cenp-C-GFP patterns were classified as aligned (3–4 discrete foci associated with the DNA), fragmented (more than four discrete foci associated with the DNA), or fragmented and misaligned (more than four discrete foci accompanied by separation of DNA from the main chromatin mass).

### Mouse strains

Study-ready aged C57BL/6J female mice (Jackson Labs, Bar Harbor USA) were used for oocyte collection at 6-12 weeks of age (young) and 40-43 weeks of age (old). Mice were housed in the Rutgers animal vivarium with constant temperature and 12-h light/12-h dark cycle. Food and water were provided *ad libitum*. All animals were maintained following the Rutgers Institutional Animal Care and Use Committee (IACUC) protocol #201702497 and in accordance with NIH policies.

### Oocyte collection and maturation

Mice were primed with 5 I.U. of pregnant mare serum gonadotrophin (PMSG) (Lee BioSolutions, Maryland Heights, MO, USA, #493-10) to induce follicle growth and recruitment and sacrificed 48 hours later. Prophase I-arrested, germinal vesicle intact oocytes were collected from antral follicles from the ovaries of female mice as previously described (Blengini and Schindler 2018). Ovaries were harvested and transferred to minimal essential medium (MEM) containing 2.5 μM milrinone (Sigma-Aldrich, St. Louis, MO, USA, #M4659) to prevent spontaneous meiotic resumption (Tsafriri et al. 1996). Follicles were punctured to release follicles and the oocytes were removed using a 100-125 μm diameter stripper pipette. Oocytes were then transferred to microdroplets of milrinone-free Chatot, Ziomek and Bavister (CZB) medium under mineral oil (Sigma-Aldrich, St. Louis, MO, USA, #M5310) and cultured at 37°C in a humidified 5% CO_2_ incubator (Chatot et al. 1989). Oocytes were cultured for 7 h to reach Metaphase I and prior to labeling of nascent translation.

### Nascent translation labeling

Translation activity in Metaphase I oocytes was assessed by detecting protein synthesis level using an Alexa Fluor 594 *L*-HPG-translation kit (ThermoFisher, Cat# C10429) as previously described (Aboelenain and Schindler 2021). Cycloheximide (Sigma-Aldrich, Cat# C7698) was added as a translation inhibitor control at 10μg/mL. Briefly, after 7 h of *in vitro* culture, oocytes were transferred into DMEM lacking methionine (Gibco, Cat # 21013-024). This culture medium also contains HPG diluted 1:50. Oocytes were incubated for 40 min, followed by fixation in 2% paraformaldehyde (PFA) in phosphate-buffered saline (PBS) for 20 min at room temperature. After fixation, oocytes were stored in blocking buffer (0.3% BSA and 0.01% Tween in PBS) overnight at 4°C. HPG signal then was detected by immunofluorescence. First, oocytes were permeabilized in PBS containing 0.2% Triton-X for 20 min and blocked again in blocking buffer for 10 min. Oocytes were then transferred into the Click-reaction solution for 40 min at room temperature in a humidified chamber, rinsed through several drops of blocking solution. After washing, Alexa Fluor 488-conjugated mouse anti-α-tubulin antibody (Invitrogen, Cat#322588) was diluted 1:100 in blocking solution and the sample was incubated for 1 h at room temperature followed by three consecutive 10 min incubations in blocking buffer. The oocytes were mounted in Vectashield (Vector Laboratories, H-1000) with 4′,6-diamidino-2-phenylindole dihydrochloride (DAPI; Life Technologies; 1:170).

### Image acquisition

Oocytes were visualized via confocal microscopy using a Leica TCS SP8 laser-scanning confocal microscope with Leica Application Suite X (version 3.5.7.23225) (Leica Microsystems). Imaging was performed with a 63X oil objective (1.40 N.A.) with a zoom setting of 2.0 or 3.5 and z-step size of 0.5 mm. Laser powers were kept constant so that HPG signal could be compared between oocytes and groups.

### Image analysis

Images were analyzed using NIH ImageJ (1.54p) software. Z-slices from each image were merged into a projection. For HPG intensity measurement, the cytoplasmic area was selected. The average intensity was recorded using the measurement tool for pixel intensity analysis. HPG signal that remained in cycloheximide-treated oocytes, was treated as background.

### Differential ribosome footprint distribution analysis

Differential ribosome footprint distribution was analyzed as previously described (Flanagan, Li, et al. 2022) using ribosome profiling data from day 2 and day 12 mature Drosophila oocytes generated by Greenblatt et al. 2019. The reference annotation was filtered to protein-coding transcripts only prior to analysis. Ribosomal P-site positions were assigned using riboWaltz, and positional ribosome footprint density profiles across coding sequences were generated using plastid. For genes with multiple annotated protein-coding transcript isoforms, a single representative transcript of matching length between day 2 and day 12 was selected per gene, choosing the isoform pair with the highest combined read count. Transcripts with coding sequences ≤200 nt or an average ribosome footprint density ≤0.15 reads per nucleotide were excluded from the analysis. Positional ribosome densities were smoothed using LOESS with a window corresponding to 5% of transcript length, and cumulative ribosome footprint distributions were calculated for each transcript. Differences in ribosome distribution between day 2 and day 12 oocytes were quantified using the Kolmogorov–Smirnov (K–S) statistic, defined as the maximum absolute distance between their cumulative ribosome footprint distributions. K–S statistics were calculated independently for each biological replicate, and the mean K–S statistic across replicates was used as the final measure of differential ribosome footprint distribution for each transcript.

## Supporting information

Supplementary Table 1: Summary of genes covered in genetic screen.

Supplementary Table 2: Genotypes for experimental flies.

Supplementary Table 3: Fly line summary for genetic screen.

## ACKNOWLEDGEMENTS

We thank the UBC Proteomics core facility for support and Nicolas Salecedo-Parros for his guidance with the ribosome purification protocol. We are grateful to Dr. Eric Jan (The University of British Columbia) for the CrPV IGR IRES plasmid, Dr. Steven DeLuca (Montana State University) for the UASz-GFP flies, and Dr. Kim McKim (Rutgers University) for the Cenp-C-GFP flies. This work is supported by the UBC Faculty of Medicine Start Up Funds, the Natural Sciences and Engineering Research Council of Canada (NSERC) and the Canada Foundation for Innovation (CFI). Mouse work was supported by the National Institutes of Health (Grant Number R35GM136340).

## DECLARATION OF INTERESTS

The authors declare no competing interests.

## DECLARATION OF AI-ASSISTED TECHNOLOGIES

Claude (Anthropic) and ChatGPT (OpenAI) were used to edit author-written text for grammar, clarity and conciseness, and to assist in writing and debugging analysis scripts for differential ribosome footprint distribution analysis. All text and code were inspected and validated by the authors. No large language model was used to generate, analyze, or interpret experimental data, or to produce any figure.

## Notes

### Competing Interest Statement

The authors have declared no competing interest.

## REFERENCES

Aboelenain, Mansour, and Karen Schindler. 2021. “Aurora Kinase B Inhibits Aurora Kinase A to Control Maternal mRNA Translation in Mouse Oocytes.” Development (Cambridge, England) 148 (21): dev199560. 10.1242/dev.199560.

Battaglia, D. E., P. Goodwin, N. A. Klein, and M. R. Soules. 1996. “Influence of Maternal Age on Meiotic Spindle Assembly in Oocytes from Naturally Cycling Women.” Human Reproduction (Oxford, England) 11 (10): 2217–22. 10.1093/oxfordjournals.humrep.a019080.

Blengini, Cecilia S., and Karen Schindler. 2018. “Immunofluorescence Technique to Detect Subcellular Structures Critical to Oocyte Maturation.” Methods in Molecular Biology (Clifton, N.J.) 1818: 67–76. 10.1007/978-1-4939-8603-3_8.

Bonus, Marissa L., Shally N. Wolf, Jessica E. Stanley, et al. 2024. “DOES RAPAMYCIN DELAY OVARIAN AGING AND DECREASE SENESCENCE? A FIRST-EVER ANALYSIS IN A NON-HUMAN PRIMATE MODEL.” Fertility and Sterility 122 (4): e344. 10.1016/j.fertnstert.2024.08.080.

Brown, James B., Nathan Boley, Robert Eisman, et al. 2014. “Diversity and Dynamics of the Drosophila Transcriptome.” Nature 512 (7515): 393–99. 10.1038/nature12962.

Cavanaugh, Kate E., M. J. Franco-Oñate, Nicole Horsley, et al. 2026. “Elevated Contractility Drives Implantation Failure in Mouse Embryos from Aged Females.” Nature Cell Biology 28 (9): 1799–813. 10.1038/s41556-026-02052-1.

Chatot, C. L., C. A. Ziomek, B. D. Bavister, J. L. Lewis, and I. Torres. 1989. “An Improved Culture Medium Supports Development of Random-Bred 1-Cell Mouse Embryos in Vitro.” Journal of Reproduction and Fertility 86 (2): 679–88. 10.1530/jrf.0.0860679.

Chiang, Teresa, Francesca E. Duncan, Karen Schindler, Richard M. Schultz, and Michael A. Lampson. 2010. “Evidence That Weakened Centromere Cohesion Is a Leading Cause of Age-Related Aneuploidy in Oocytes.” Current Biology: CB 20 (17): 1522–28. 10.1016/j.cub.2010.06.069.

Danielson, Katie J., Kayla L. Judson, and Ethan J. Greenblatt. 2024. Reproductive Ageing: Declining Translational Capacity as a Potential Driver for Oocyte Meiotic Instability. Reproduction. November 1. 10.1530/REP-24-0198.

De Leon, Victor, Andrew Johnson, and Rosemary Bachvarova. 1983. “Half-Lives and Relative Amounts of Stored and Polysomal Ribosomes and Poly(A)+ RNA in Mouse Oocytes.” Developmental Biology 98 (2): 400–408. 10.1016/0012-1606(83)90369-X.

Del Llano, Edgar, Tomas Masek, Lenka Gahurova, et al. 2020. “Age-related Differences in the Translational Landscape of Mammalian Oocytes.” Aging Cell 19 (10): e13231. 10.1111/acel.13231.

Dudko, Nataliia, Tereza Ilcikova, Natalie Novotna, et al. 2026. “Oocyte Age-Dependent DNA Damage Can Be Reverted by the DNA Repair Competent Karyoplasm of Young Oocytes.” Aging Cell 25 (1): e70300. 10.1111/acel.70300.

Flanagan, Keegan, Alireza Baradaran-Heravi, Qi Yin, Khanh Dao Duc, Allan C. Spradling, and Ethan J. Greenblatt. 2022. “FMRP-Dependent Production of Large Dosage-Sensitive Proteins Is Highly Conserved.” Genetics 221 (4): iyac094. 10.1093/genetics/iyac094.

Flanagan, Keegan, Wanxin Li, Ethan J. Greenblatt, and Khanh Dao Duc. 2022. “End-to-End Pipeline for Differential Analysis of Pausing in Ribosome Profiling Data.” STAR Protocols 3 (3): 101605. 10.1016/j.xpro.2022.101605.

Graveley, Brenton R., Angela N. Brooks, Joseph W. Carlson, et al. 2011. “The Developmental Transcriptome of Drosophila Melanogaster.” Nature 471 (7339): 473–79. 10.1038/nature09715.

Greenblatt, Ethan J., Rebecca Obniski, Claire Mical, and Allan C. Spradling. 2019. “Prolonged Ovarian Storage of Mature Drosophila Oocytes Dramatically Increases Meiotic Spindle Instability.” eLife 8 (November): e49455. 10.7554/eLife.49455.

Gruhn, Jennifer R., Agata P. Zielinska, Vallari Shukla, et al. 2019. “Chromosome Errors in Human Eggs Shape Natural Fertility over Reproductive Life Span.” Science 365 (6460): 1466–69. 10.1126/science.aav7321.

Holubcova, Zuzana, Blayney Martyn, Elder Kay, and Schuh Melina. 2015. “Error-Prone Chromosome-Mediated Spindle Assembly Favors Chromosome Segregation Defects in Human.” Science 348 (6239): 1143–47. 10.1126/science.aaa4788.

Huang, Jiana, Peigen Chen, Lei Jia, et al. 2023. “Multi-Omics Analysis Reveals Translational Landscapes and Regulations in Mouse and Human Oocyte Aging.” Advanced Science 10 (26): 2301538. 10.1002/advs.202301538.

Hughes, Christopher S., Sophie Moggridge, Torsten Müller, Poul H. Sorensen, Gregg B. Morin, and Jeroen Krijgsveld. 2019. “Single-Pot, Solid-Phase-Enhanced Sample Preparation for Proteomics Experiments.” Nature Protocols 14 (1): 68–85. 10.1038/s41596-018-0082-x.

Khajuria, Rajiv K., Mathias Munschauer, Jacob C. Ulirsch, et al. 2018. “Ribosome Levels Selectively Regulate Translation and Lineage Commitment in Human Hematopoiesis.” Cell 173 (1): 90–103.e19. 10.1016/j.cell.2018.02.036.

Kim, Harper S., and Andrew M. Pickering. 2023. “Protein Translation Paradox: Implications in Translational Regulation of Aging.” Frontiers in Cell and Developmental Biology 11 (January). 10.3389/fcell.2023.1129281.

Lagirand-Cantaloube, Julie, Cendrine Ciabrini, Sophie Charrasse, et al. 2017. “Loss of Centromere Cohesion in Aneuploid Human Oocytes Correlates with Decreased Kinetochore Localization of the Sac Proteins Bub1 and Bubr1.” Scientific Reports 7 (1): 44001. 10.1038/srep44001.

Lebaron, Simon, Claudia Schneider, Robert W. Van Nues, et al. 2012. “Proofreading of Pre-40S Ribosome Maturation by a Translation Initiation Factor and 60S Subunits.” Nature Structural & Molecular Biology 19 (8): 744–53. 10.1038/nsmb.2308.

Leblanc, Thierry, Laurence Faivre, Jorg Meerpohl, et al. 2005. “High Risk Pregnancies in Women with Diamond Blackfan Anaemia: A Survey of 64 Pregnancies from the French and German Registries.” Blood 106 (11): 3549. 10.1182/blood.V106.11.3549.3549.

Lister, Lisa Martine, Anna Kouznetsova, Louise Ann Hyslop, et al. 2010. “Age-Related Meiotic Segregation Errors in Mammalian Oocytes Are Preceded by Depletion of Cohesin and Sgo2.” Current Biology 20 (17): 1511–21. 10.1016/j.cub.2010.08.023.

Marygold, Steven J., John Roote, Gunter Reuter, et al. 2007. “The Ribosomal Protein Genes and Minute Loci of Drosophila Melanogaster.” Genome Biology 8 (10): R216. 10.1186/gb-2007-8-10-r216.

Nakagawa, Shoma, and Greg FitzHarris. 2017. “Intrinsically Defective Microtubule Dynamics Contribute to Age-Related Chromosome Segregation Errors in Mouse Oocyte Meiosis-I.” Current Biology: CB 27 (7): 1040–47. 10.1016/j.cub.2017.02.025.

Natsidis, Paschalis, Philipp H. Schiffer, Irepan Salvador-Martínez, and Maximilian J. Telford. 2019. “Computational Discovery of Hidden Breaks in 28S Ribosomal RNAs across Eukaryotes and Consequences for RNA Integrity Numbers.” Scientific Reports 9 (1): 19477. 10.1038/s41598-019-55573-1.

Perkins, Adrienne T., Thomas M. Das, Lauren C. Panzera, and Sharon E. Bickel. 2016. “Oxidative Stress in Oocytes during Midprophase Induces Premature Loss of Cohesion and Chromosome Segregation Errors.” Proceedings of the National Academy of Sciences of the United States of America 113 (44): E6823–30. 10.1073/pnas.1612047113.

Petrova, Boryana, Keke Liu, Caiping Tian, et al. 2018. “Dynamic Redox Balance Directs the Oocyte-to-Embryo Transition via Developmentally Controlled Reactive Cysteine Changes.” Proceedings of the National Academy of Sciences 115 (34): E7978–86. 10.1073/pnas.1807918115.

Rappsilber, Juri, Yasushi Ishihama, and Matthias Mann. 2003. “Stop and Go Extraction Tips for Matrix-Assisted Laser Desorption/Ionization, Nanoelectrospray, and LC/MS Sample Pretreatment in Proteomics.” Analytical Chemistry 75 (3): 663–70. 10.1021/ac026117i.

Robledo, Sara, Rachel A. Idol, Dan L. Crimmins, Jack H. Ladenson, Philip J. Mason, and Monica Bessler. 2008. “The Role of Human Ribosomal Proteins in the Maturation of rRNA and Ribosome Production.” RNA 14 (9): 1918–29. 10.1261/rna.1132008.

Romasko, Edward J., Dasari Amarnath, Uros Midic, and Keith E. Latham. 2013. “Association of Maternal mRNA and Phosphorylated EIF4EBP1 Variants With the Spindle in Mouse Oocytes: Localized Translational Control Supporting Female Meiosis in Mammals.” Genetics 195 (2): 349–58. 10.1534/genetics.113.154005.

Sakakibara, Yogo, Shu Hashimoto, Yoshiharu Nakaoka, Anna Kouznetsova, Christer Höög, and Tomoya S. Kitajima. 2015. “Bivalent Separation into Univalents Precedes Age-Related Meiosis I Errors in Oocytes.” Nature Communications 6 (1): 7550. 10.1038/ncomms8550.

Severance, Ashley L., and Keith E. Latham. 2017. “PLK1 Regulates Spindle Association of Phosphorylated Eukaryotic Translation Initiation Factor 4E-Binding Protein and Spindle Function in Mouse Oocytes.” American Journal of Physiology - Cell Physiology 313 (5): C501–15. 10.1152/ajpcell.00075.2017.

Shcherbik, Natalia, and Dimitri G. Pestov. 2019. “The Impact of Oxidative Stress on Ribosomes: From Injury to Regulation.” Cells 8 (11): 1379. 10.3390/cells8111379.

Sobek, Ales, Emil Tkadlec, Eva Klaskova, and Martin Prochazka. 2020. “Cytoplasmic Transfer Improves Human Egg Fertilization and Embryo Quality: An Evaluation of Sibling Oocytes in Women with Low Oocyte Quality.” Reproductive Sciences 28 (5): 1362–69. 10.1007/s43032-020-00371-8.

Sousa Martins, Joao P., Xueqing Liu, Ashwini Oke, et al. 2016. “DAZL and CPEB1 Regulate mRNA Translation Synergistically during Oocyte Maturation.” Journal of Cell Science 129 (6): 1271–82. 10.1242/jcs.179218.

Stein, Kevin C., Fabián Morales-Polanco, Joris van der Lienden, T. Kelly Rainbolt, and Judith Frydman. 2022. “Ageing Exacerbates Ribosome Pausing to Disrupt Cotranslational Proteostasis.” Nature 601 (7894): 637–42. 10.1038/s41586-021-04295-4.

Susor, Andrej, Denisa Jansova, Renata Cerna, et al. 2015. “Temporal and Spatial Regulation of Translation in the Mammalian Oocyte via the mTOR–eIF4F Pathway.” Nature Communications 6 (1): 6078. 10.1038/ncomms7078.

Takahashi, Nozomi, Federica Franciosi, Enrico Maria Daldello, et al. 2023. “CPEB1-Dependent Disruption of the mRNA Translation Program in Oocytes during Maternal Aging.” Nature Communications 14 (1): 416. 10.1038/s41467-023-35994-3.

Tsafriri, A., Sang-Young Chun, Ruobo Zhang, A. J. W. Hsueh, and M. Conti. 1996. “Oocyte Maturation Involves Compartmentalization and Opposing Changes of cAMP Levels in Follicular Somatic and Germ Cells: Studies Using Selective Phosphodiesterase Inhibitors.” Developmental Biology 178 (2): 393–402. 10.1006/dbio.1996.0226.

Tsutsumi, Makiko, Reiko Fujiwara, Haruki Nishizawa, et al. 2014. “Age-Related Decrease of Meiotic Cohesins in Human Oocytes.” PLOS ONE 9 (5): e96710. 10.1371/journal.pone.0096710.

Tufano, Andrea M., Diana S. Osorio, Eva Atsidaftos, Cristina P. Sison, Jeffrey M. Lipton, and Adrianna Vlachos. 2014. “Deleterious Consequences of Diamond Blackfan Anemia on Reproductive Health and Pregnancy Outcomes: A Report from the Diamond Blackfan Anemia Registry (DBAR).” Blood 124 (21): 4399–4399. 10.1182/blood.V124.21.4399.4399.

Yin, Lin, Aleksandar I. Mihajlović, Gongshe Yang, and Greg FitzHarris. 2023. “Kinetochore Deterioration Concommitant with Centromere Weakening during Aging in Mouse Oocyte Meiosis-I.” The FASEB Journal 37 (5): e22922. 10.1096/fj.202300062R.

Yun, Yan, Janet E. Holt, Simon Lane, Eileen McLaughlin, Julie Merriman, and Keith Jones. 2014. “Reduced Ability to Recover from Spindle Disruption and Loss of Kinetochore Spindle Assembly Checkpoint Proteins in Oocytes from Aged Mice.” Cell Cycle 13 (12): 1938–47. 10.4161/cc.28897.

Zhang, Yu, Jie Bai, Na Li, and Bo Xiong. 2026. “Insufficiency of Cohesin Regulatory Protein PDS5B Triggers Meiotic Failure and Aneuploidy in Oocytes during Advanced Maternal Aging.” Proceedings of the National Academy of Sciences 123 (31): e2605947123. 10.1073/pnas.2605947123.

Zielinska, Agata P., Eirini Bellou, Ninadini Sharma, et al. 2019. “Meiotic Kinetochores Fragment into Multiple Lobes upon Cohesin Loss in Aging Eggs.” Current Biology 29 (22): 3749–3765.e7. 10.1016/j.cub.2019.09.006.

Zubradt, Meghan, Paromita Gupta, Sitara Persad, Alan M. Lambowitz, Jonathan S. Weissman, and Silvi Rouskin. 2017. “DMS-MaPseq for Genome-Wide or Targeted RNA Structure Probing in Vivo.” Nature Methods 14 (1): 1. 10.1038/nmeth.4057.

