## Supplemental Materials for "Declining ribosome function limits oocyte longevity"

**TABLE LEGENDS**

Supplementary Table 1: Summary of genes covered in genetic screen.

Supplementary Table 2: Genotypes for experimental flies.

Supplementary Table 3: Fly line summary for genetic screen.

**FIGURES**

**Figure S1**

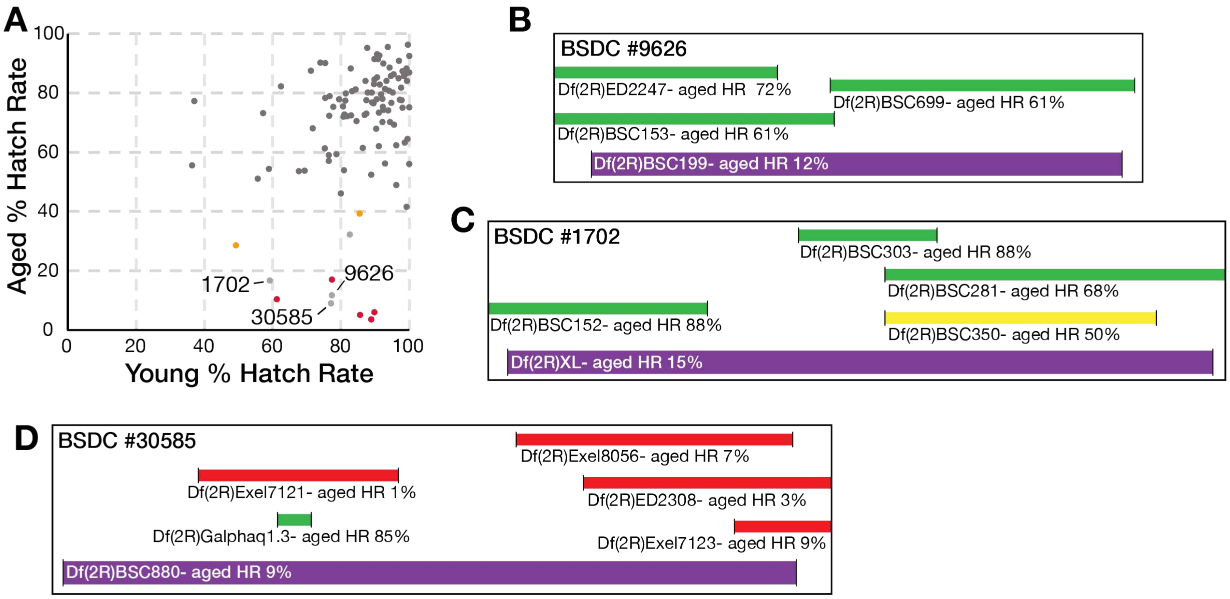


**Figure S1. Genomic regions from overlapping deficiency lines lacking a single candidate gene. (A)** Summary of screen results, each point representing a single deficiency line; red and yellow indicate strong and moderate hits, respectively. Light grey points denote deficiency lines for which no single candidate gene could be isolated; each is labeled with its corresponding Bloomington Stock Center Deficiency (BSDC) identification number. **(B–D)** Approximate genomic positions of the deleted region for each indicated deficiency line (purple) and its overlapping deficiency lines (above).

**Figure S2**

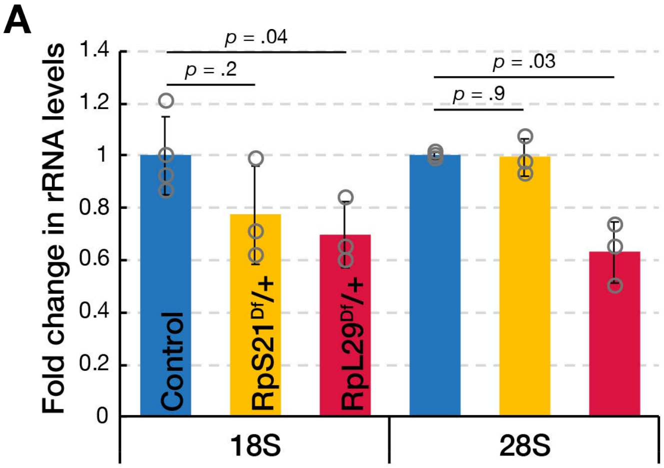


**Figure S2. RpL29 reduction decreases both 18S and 28S rRNA, while RpS21 reduction selectively affects 18S rRNA. (A)** rRNA levels in *RpL29*$^{Df/}$+ and *RpS21*$^{Df}$/+ were quantified by Bioanalyzer trace. *RpL29*$^{Df/}$+ showed significantly reduced levels of both 18S and 28S rRNA, whereas *RpS21*$^{Df}$/+ showed no change in 28S rRNA and a non-significant trend toward reduced 18S rRNA. Welch's unpaired *t*-test was used for statistical comparisons.

**Figure S3**

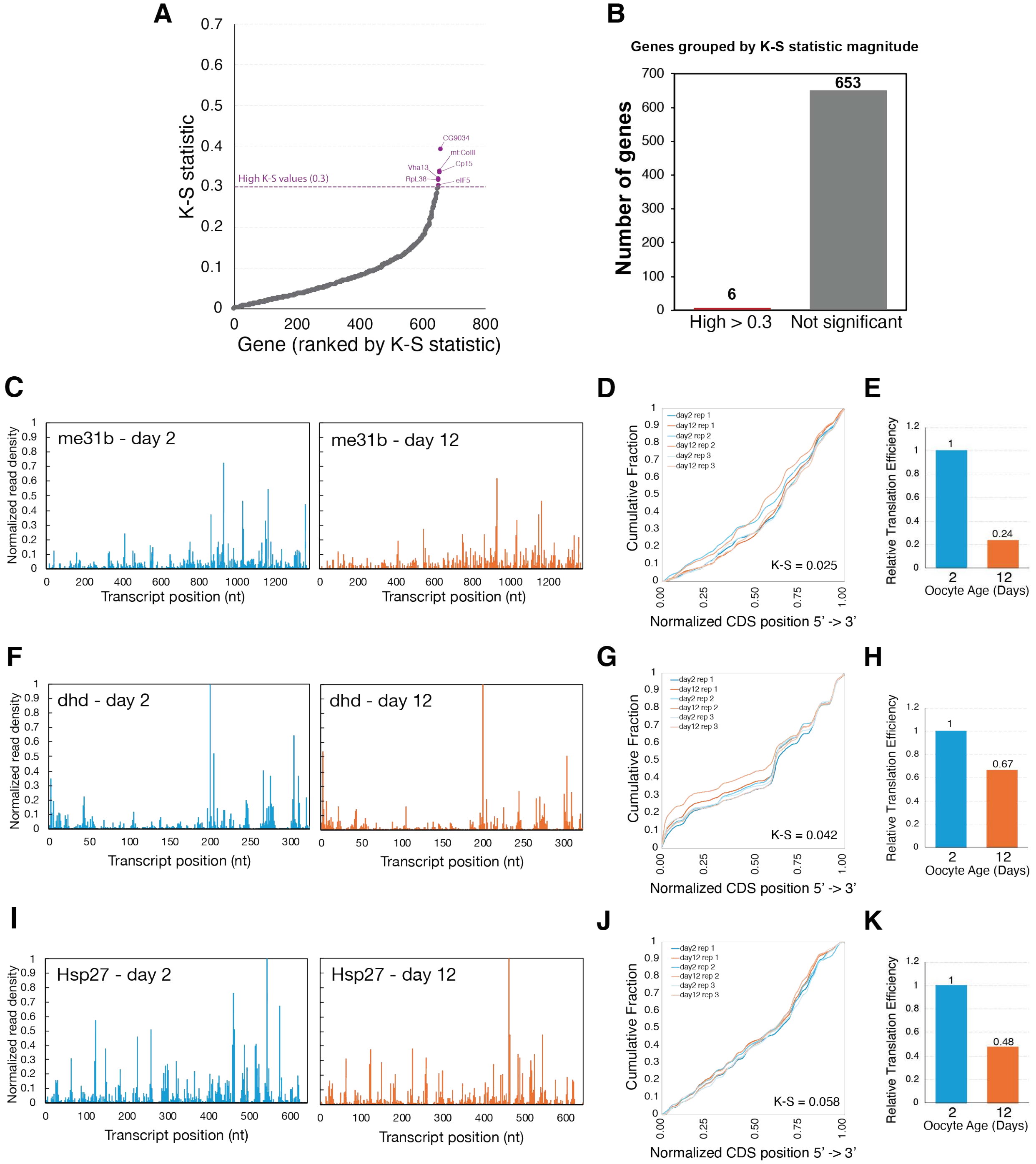


**Figure S3. Ribosome footprint distribution is largely unchanged during aging. (A)** Kolmogorov-Smirnov (K-S) statistics quantifying the shift in cumulative ribosome footprint distribution between young (day 2) and old (day 12) oocytes, calculated for 659 genes and ranked from lowest to highest. The K-S statistic threshold for a significant distributional shift (0.3) was defined by simulation-based modeling from (Flanagan, Baradaran-Heravi, et al. 2022).

**(B)** Genes were classified as high (K-S ≥ 0.3) or not significant (K-S < 0.3). A high K-S statistic indicates a transcript with altered ribosome footprint distribution with age, while a low K-S statistic indicates a distribution that remains highly similar between young and old oocytes. **(C–E)** Representative example for *me31B*: **(C)** Ribosome read density along the transcript in young (blue) and old (orange) oocytes; **(D)** Cumulative footprint distribution across replicates, showing minimal change with age (K-S = 0.025); **(E)** Relative translation efficiency (TE) calculated with ribosome profiling data from (Greenblatt et al. 2019) in day 2 versus day 12 oocytes, showing reduced TE despite an unchanged footprint distribution. **(F–H)** As in (C–E), for the transcript *dhd* (K-S = 0.042). **(I–K)** As in (C–E), for the transcript *Hsp27* (K-S = 0.058).

**Figure S4**

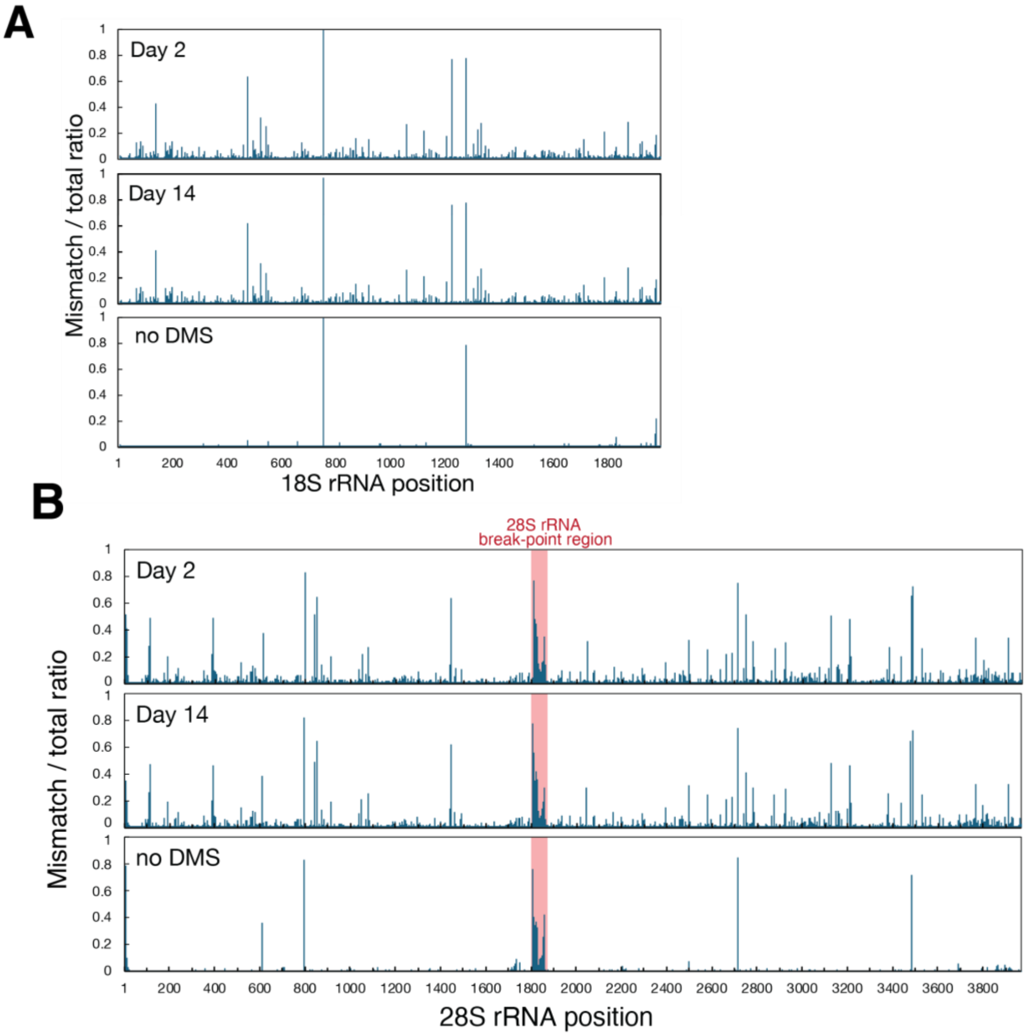


**Figure S4. Specificity of DMS-MapSeq profiling (A)** Raw mismatch/total ratios for 18S rRNA in the untreated (no DMS) control versus the *in vivo* DMS-treated sample (50% v/v DMS, 5 min), plotted for all nucleotides (A, C, G, U). The untreated control shows low background mutation rates across the transcript. Positions with elevated mutation ratios in the untreated control were excluded from downstream analysis of the DMS-treated sample. **(B)** Raw mismatch/total ratios for 28S rRNA, as in (A). The red shaded region marks the 28S hidden break, a site of physical rRNA backbone cleavage that causes a local drop in sequencing coverage and a correspondingly inflated mismatch/total ratio, independent of DMS treatment. Positions with elevated mutation ratios in the untreated control, including the hidden break region, were excluded from downstream analysis of the DMS-treated sample. See Figure 4C and 4E for filtered reactivity profiles used in downstream analysis.

**Figure S5**

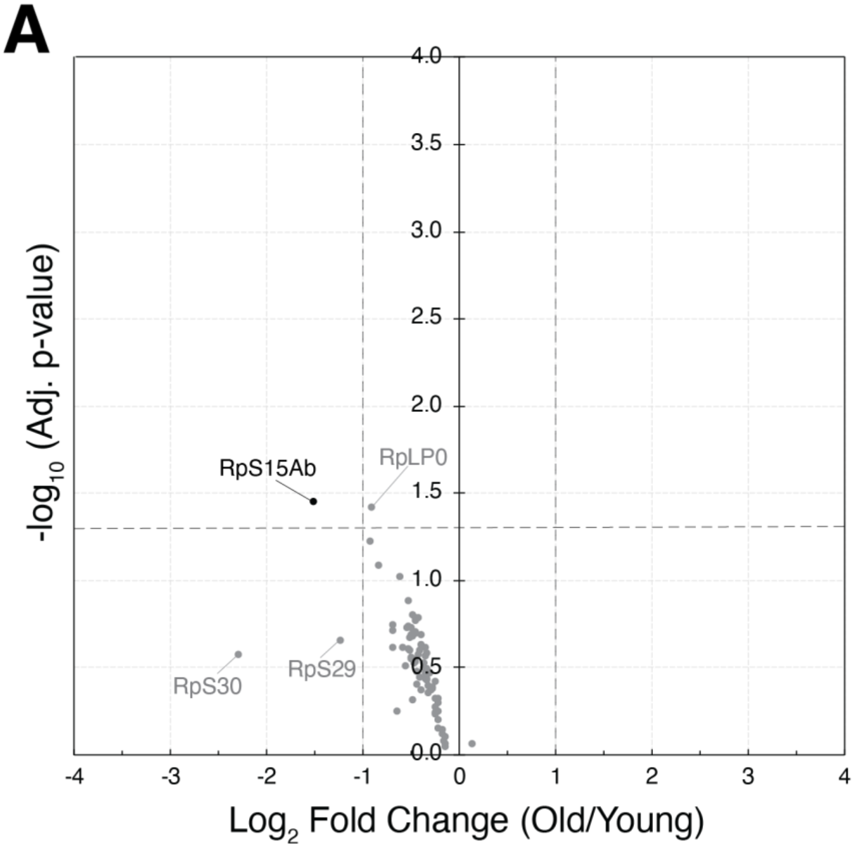


**Figure S5. Ribosomal protein composition is largely stable during aging, with the exception of RpS15Ab. (A)** Volcano plot of mass spectrometry data from pelleted ribosome fractions showing that RpS15Ab is significantly decreased with age, while overall ribosomal protein stoichiometry remains largely unchanged, consistent with Figure 4G. Data are the same as presented in Figure 4G, replotted to highlight ribosomal protein abundance. Differential abundance was assessed using Welch's two-sample *t*-test with Benjamini-Hochberg FDR correction; dashed lines indicate thresholds of FDR < 0.05 and $|\log_2$ fold change$| > 1$. N = 4 biological replicates per condition.

**Figure S6**

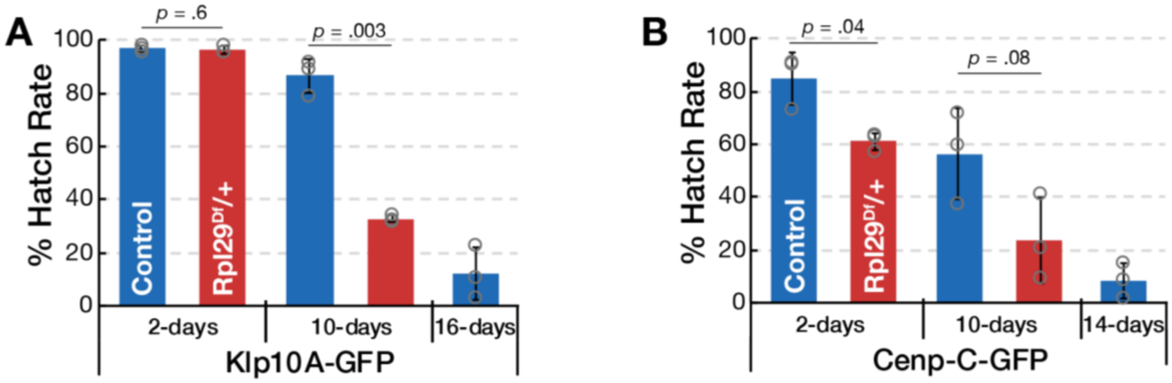


**Figure S6. RpL29 haploinsufficiency accelerates age-related oocyte viability decline. (A)** In the Klp10A-GFP background, hatch rate of RpL29$^{Df}$/+ oocytes shows an accelerated decline with age, with no difference in young (2-day-old) oocytes. **(B)** In the Cenp-C-GFP background, hatch rate of RpL29$^{Df}$/+ oocytes is reduced at 2 days old; a similar trend is observed at day 10 but does not reach statistical significance. Welch's unpaired *t*-test was used for statistical comparisons.
